# SpaceMACE: Spatially-Aware Tumour Cell Annotation for Spatial Transcriptomics

**DOI:** 10.64898/2026.11.16.763092

**Authors:** Davide D’Agostini, Tamim Abdelaal

## Abstract

Accurate discrimination between tumour and non-tumour cells is essential for interpreting spatial transcriptomics data and identifying spatially resolved therapeutic targets. Existing tumour-cell annotation approaches were primarily developed for single-cell RNA sequencing and do not explicitly incorporate the spatial organization of cells.

Here, we introduce SpaceMACE, an extension of the MACE tumour-cell annotation framework for sequencing-based and imaging-based spatial transcriptomics. SpaceMACE first combines reference-based cell-type classification with copy-number-variation-based tumour classification. The resulting consensus annotations are subsequently refined using spatial proximity and gene-expression similarity.

We evaluated SpaceMACE across multiple spatial transcriptomics platforms, including Stereo-seq, Visium HD, CosMx and Xenium, and across colorectal, hepatocellular and ovarian cancer datasets. SpaceMACE retained the high precision of the original MACE framework while improving the spatial coherence of tumour-cell annotations. Together, these results demonstrate that SpaceMACE provides a scalable and platform-independent framework for tumour-cell annotation in spatial transcriptomics and supports the spatial investigation of tumour-specific molecular targets.

## 1 Introduction

Targeted cancer therapies derive most of their benefit from tumour-specific drug activity, minimising off-target effects while maximising efficacy in malignant tissue [1, 2]. To discover robust tumour-specific targets, we must first delineate which cells in a specimen are truly tumour and which belong to the surrounding tumour micro-environment (TME). Single-cell RNA sequencing (scRNA-seq) has transformed cancer biology by resolving heterogeneous tissue into transcriptionally defined cell states and lineages [3, 4]. In tumours, scRNA-seq enables the identification of malignant epithelial programs and sub-clones, mapping of immune composition and activation states, and characterization of stromal elements such as cancer-associated fibroblasts and endothelial cells [5–8]. Despite its power, reliable tumour/normal separation is non-trivial: dissociation can bias cell recovery, sparsity and batch effects complicate modelling, and malignant cells may transcriptionally resemble non-malignant epithelial cells [9–11]. These challenges motivate annotation strategies that combine complementary signals and emphasize precision to avoid false tumour calls that could mislead target selection.

Within automated annotators for cancer scRNA-seq, two complementary families dominate. Reference-based classifiers, such as scATOMIC [12], learn from curated cancer references and hierarchical cell-type ontologies to label new cells from expression patterns; they excel for well-represented phenotypes, but can conflate malignant and normal epithelium when their gene-expression profiles are similar. CNV-based methods, such as inferCNV [13], copyKAT [14] or SCEVAN [15], infer large-scale copy number alterations from expression, providing an orthogonal genomic signal characteristic of many cancers; however, CNV evidence can be noisy at low coverage or less informative in tumours with subtle aberrations. To capitalise on their strengths while mitigating their weaknesses, MACE [16] applies a conservative consensus: a cell is called a tumour only when scATOMIC and SCEVAN agree. This consensus deliberately prioritises precision and reduces false positives in various cancer types, providing a stable baseline for downstream target discovery.

Although scRNA-seq profiles dissociated cells, spatial transcriptomics (ST) preserves tissue architecture, allowing direct study of tumour-stroma interfaces, immune niches, and intratumour heterogeneity in situ [17–21]. Two broad ST modalities can be defined: sequencing-based ST (sST) measures whole-transcriptome gene sets on spatially barcoded arrays; it offers broad gene coverage but variable effective resolution and spot/pixel sparsity, as seen in platforms such as Visium HD, Stereo-seq, and Open-ST [22–24]. Imaging-based ST (iST), such as Xenium and CosMx, quantifies preselected gene panels at near single-cell resolution; it provides precise localisation but limited gene coverage [25]. Accurate annotation of tumour cells is especially important in the spatial context. Because ST preserves the physical positions of cells within tissue, misclassifying malignant and normal cells does not just affect cell-type counts, it directly distorts our picture of how tumours are spatially organised. For example, incorrectly labelling normal cells as tumour inflates apparent tumour density in a region, confounds analyses of how cancer and immune or stromal cells communicate, and can create or obscure gene expression patterns that would otherwise guide target discovery. As ST is increasingly used to map where distinct malignant subpopulations reside within a tumour, knowing which cells are truly malignant becomes a prerequisite for any meaningful spatial analysis. Bringing the same precision-focused annotation strategies developed for scRNA-seq to the spatial domain is therefore an important step toward reliable and spatially resolved cancer biology [26, 27].

Despite these opportunities, working with ST data presents substantial computational and analytical challenges that do not arise in scRNA-seq. Spatial datasets are orders of magnitude larger: a single Visium HD slide operates at 2*µ*m pixel resolution or stereo-seq that operates at 0.5*µ*m, yielding tens of millions of raw capture spots, while imaging-based platforms such as Xenium and CosMx produce hundreds of thousands of single-cell measurements per slide. This scale strains conventional single-cell tooling, which was not designed for cell counts in this range. Beyond size, each ST modality introduces its own data-specific challenges: sequencing-based platforms suffer from severe per-pixel sparsity that requires aggregation into larger units before meaningful inference is possible, while imaging-based platforms are limited to targeted gene panels of a few thousand genes, making tools that depend on whole-transcriptome signals inapplicable out of the box. Several methods have been developed to address aspects of ST data analysis in cancer: SlideCNA [27] extends CNV-based tumour detection to spatial data, SpaCET [28] deconvolves the tumour and immune cell fractions from ST spots, and Cancer-Finder [26] annotates malignant spots using a domain generalisation approach. However, none of these provides an end-to-end precision-focused tumour annotation pipeline that operates across both sST and iST modalities while preserving the spatial context of every cell. This gap motivates the present work.

Building on MACE’s consensus framework for scRNA-seq, our goal is to extend MACE to spatial transcriptomics by incorporating spatial neighbourhood information into the labelling pipeline. For sST, we stabilise expression by aggregating pixels into biologically informed spatial units using ENACT [29]; for iST, we recover the coverage of the whole-transcriptome by aligning the data from the targeted panel with the scRNA-seq using SpaGE [30]. To handle the scale of spatial datasets, SCEVAN was modified to replace its internal clustering routine with a memory-efficient alternative, removing a hard cell-count ceiling that would otherwise prevent the processing of large spatial cohorts. We then apply MACE and incorporate spatial refinement through two complementary models: SpaceMACE-S, which enforces label consistency using spatial proximity alone, and SpaceMACE, which combines spatial and transcriptomics similarity for a richer refinement. The overarching objective is to produce spatially coherent high-precision tumour annotations in situ that better support tumour-specific target discovery. The resulting SpaceMACE framework is applicable across both sST and iST modalities and fully leverages ST data by jointly exploiting gene expression and the spatial localisation of cells to achieve reliable tumour versus normal separation.

## 2 Methods

### 2.1 Tumour cell identification methods

#### 2.1.1 MACE

MACE combines a reference-based classifier (scATOMIC) with a copy-number variation (CNV)-based method (SCEVAN) and assigns final labels through a consensus strategy. First, scATOMIC is applied to the entire scRNA-seq matrix to produce a set of high-level tumour candidates (epithelial lineage) and flag low-quality cells. SCE-VAN is then applied exclusively to the scATOMIC epithelial/tumour candidate and undefined subsets, to separate malignant from normal epithelium. By default, SCE-VAN receives a diploid reference of 50 cells randomly sampled from the scATOMIC *normal* set, replacing its internal normal-finding step; this stabilises the CNV base-line and reduces runtime without altering the decision logic. The final cell labels are assigned as follows:

- **tumour:** both scATOMIC and SCEVAN independently call the cell a tumour.
- **Normal epithelial:** scATOMIC calls tumour, but SCEVAN calls non-tumour.
- **Normal:** scATOMIC calls non-tumour (SCEVAN is not applied).
- **Filtered:** flagged as low quality by scATOMIC or SCEVAN.

All method defaults are used unless stated otherwise; scATOMIC is run without fine-grained options and SCEVAN via its standard CNA pipeline. Implementation follows the public MACE R package (Seurat-compatible wrappers). This consensus design prioritises precision and reduces false positives between cancer types compared to running scATOMIC or SCEVAN alone. To support the larger cell counts encountered in the iST and sST datasets, SCEVAN required a targeted modification. The original implementation relies on stats::hclust for hierarchical clustering of the CNV profiles, which imposes an internal limit that causes the method to fail when the tumour compartment exceeds *∼* 65,000 cells. We resolved this by replacing stats::hclust with fastcluster::hclust [31], a drop-in replacement that lifts this constraint through a more memory-efficient implementation of the same agglomerative clustering algorithm. This substitution does not alter the underlying CNV calling logic or any decision thresholds.

#### 2.1.2 SpaceMACE

SpaceMACE, fuses spatial and gene-expression information. To evaluate the contribution of gene expression, we also tested a spatial-only variant, SpaceMACE-S, obtained by setting *α* = 1 in Equation 2, which reduces the fused weight to the Gaussian spatial kernel alone. Both variants share the same reassignment rule and protected-label handling.

#### Shared reassignment rule

For each cell *i*, the *k* nearest neighbours were identified in physical (*x, y*) coordinates using Euclidean distance. Filtered cells were excluded from neighbour queries and were not reassigned. A weighted vote was computed over the valid neighbours. Cell *I* was reassigned to the label with the maximum aggregate weight if the corresponding weight fraction was at least *τ* :

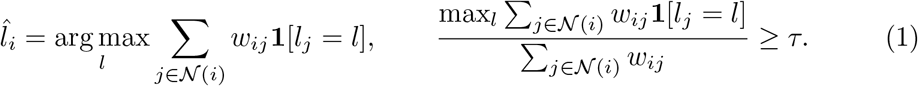

Here, *N* (*i*) denotes the valid spatial neighbourhood of cell *i, l*_*j*_ denotes the label of neighbour *j*, and *w*_*ij*_ denotes the variant-specific neighbour weight defined below.

#### SpaceMACE

SpaceMACE computes neighbour weights as a convex combination of spatial and gene-expression similarity:

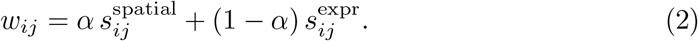

Spatial similarity was defined by a Gaussian kernel,

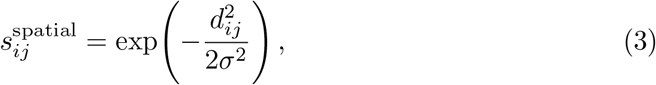

where *d*_*ij*_ is the Euclidean distance between cells *i* and *j*. The bandwidth *σ* was set to the median distance to the *k*-th nearest neighbour across all valid cells. Expression similarity was computed as cosine similarity between log-normalised expression vectors and rescaled to [0, 1]:

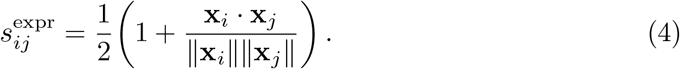

The parameters were *k* = 30, *α* = 0.4, and *τ* = 0.7.

#### SpaceMACE-S (spatial-only ablation)

To isolate the contribution of gene expression, we tested a spatial-only variant obtained by setting *α* = 1 in Equation 2, which reduces the neighbour weight to the Gaussian kernel alone:

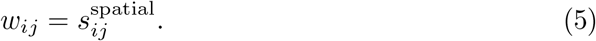

All other settings were kept identical, with *k* = 30 and *τ* = 0.6.

### 2.2 Datasets

To evaluate the performance of SpaceMACE across different spatial transcriptomics technologies, we used matched multi-indication datasets from the SPATCH project [32]. SPATCH provides paired ST and scRNA-seq datasets from the same tumour specimens together with author-provided cell-type annotations, enabling systematic evaluation of tumour-cell classification across multiple cancer types. Three cancer indications were included throughout this study: colorectal adenocarcinoma (COAD), hepatocellular carcinoma (HCC), and ovarian cancer (OV).

The spatial datasets span both major classes of ST technologies. Sequencing-based spatial transcriptomics (sST), represented by Visium HD and Stereo-seq, provides near whole-transcriptome measurements but suffers from sparse transcript capture at very high spatial resolution. Imaging-based spatial transcriptomics (iST), represented by Xenium and CosMx, provides single-cell or near-single-cell resolution through targeted RNA panels, offering precise cellular localisation at the expense of transcriptome coverage. The matched scRNA-seq datasets serve both as a reference for SpaGE transcriptome imputation and as a reference for scRNA-seq-guided bin-size selection in sequencing-based platforms.

#### 2.2.1 Sequencing-based spatial transcriptomics

Visium HD captures transcriptomes on a spatially barcoded array with approximately 2 *µ*m resolution [33]. The extremely high spatial sampling results in sparse individual capture bins that require aggregation before downstream analysis. Three representations were evaluated: the original 2 *µ*m bins, the vendor-provided 8 *µ*m supercells, and ENACT-derived segmentation-based cells.

Stereo-seq uses DNA nanoball arrays to generate whole-transcriptome measurements at approximately 0.5 *µ*m spatial resolution [34]. Similar to Visium HD, neighbouring pixels were aggregated before downstream analysis to obtain sufficient transcript counts for tumour-cell classification. Table 1 summarises the dimensions of the sequencing-based datasets.

**Table 1.**
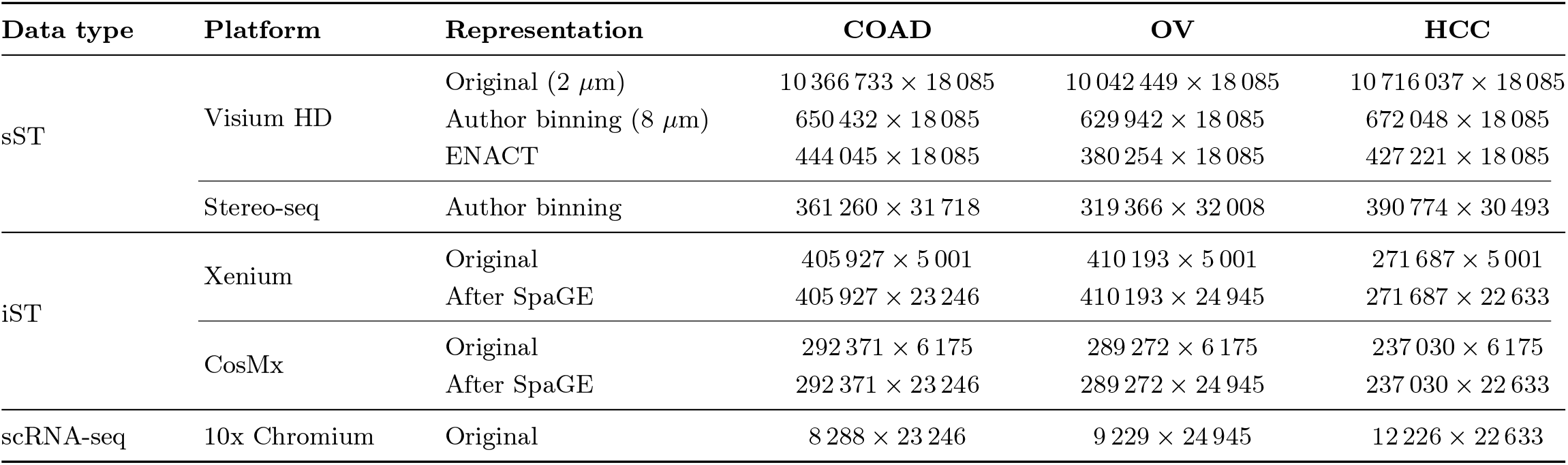
Dimensions of the spatial transcriptomics and matched scRNA-seq datasets. Values are reported as number of observations *×* number of genes.

| Data type | Platform | Representation | COAD | OV | HCC |
|---|---|---|---|---|---|
| sST | Visium HD | Original (2 μm) | 10 366 733 × 18 085 | 10 042 449 × 18 085 | 10 716 037 × 18 085 |
|  |  | Author binning (8 μm) | 650 432 × 18 085 | 629 942 × 18 085 | 672 048 × 18 085 |
|  |  | ENACT | 444 045 × 18 085 | 380 254 × 18 085 | 427 221 × 18 085 |
|  | Stereo-seq | Author binning | 361 260 × 31 718 | 319 366 × 32 008 | 390 774 × 30 493 |
| iST | Xenium | Original | 405 927 × 5 001 | 410 193 × 5 001 | 271 687 × 5 001 |
|  |  | After SpaGE | 405 927 × 23 246 | 410 193 × 24 945 | 271 687 × 22 633 |
|  | CosMx | Original | 292 371 × 6 175 | 289 272 × 6 175 | 237 030 × 6 175 |
|  |  | After SpaGE | 292 371 × 23 246 | 289 272 × 24 945 | 237 030 × 22 633 |
| scRNA-seq | 10x Chromium | Original | 8 288 × 23 246 | 9 229 × 24 945 | 12 226 × 22 633 |

#### 2.2.2 Imaging-based spatial transcriptomics

Xenium and CosMx measure targeted RNA panels directly within intact tissue using multiplexed imaging approaches [35, 36]. Both platforms provide single-cell or near-single-cell resolution through vendor-generated cell segmentations but measure only a subset of the transcriptome. To enable copy-number inference with MACE, transcriptome-wide expression profiles were predicted using SpaGE. Table 1 summarises the dimensions of the imaging-based datasets before and after SpaGE imputation.

#### 2.2.3 Reference single-cell RNA sequencing

Matched scRNA-seq datasets were generated from dissociated primary tumour tissue originating from the same COAD, OV, and HCC specimens as the spatial datasets [32]. Samples with at least 85% viability were processed using the 10x Genomics Chromium Single Cell 3’v3.1 workflow. These datasets were used as references for SpaGE transcriptome imputation and for scRNA-seq-guided bin-size selection. Table 1 summarises the dimensions of the scRNA-seq datasets.

#### 2.2.4 ENACT binning

ENACT is a computational library designed for the analysis of Visium HD spatial transcriptomics data [29]. It takes as input the raw Visium HD dataset at 2 *µ*m pixel/spot resolution, the tissue staining image (haematoxylin and Eosin (H&E) or Immunofluorescence (IF)), and a parquet file containing spatial coordinates and intissue classification metadata, which together are used to define cell boundaries and dimensions.

Cell segmentation is performed using StarDist [37], a deep learning-based nuclear detection algorithm. In this study, the default pre-defined hyperparameters provided by ENACT were used without modification.

Since Visium HD operates at 2 *µ*m resolution, individual bins frequently overlap with multiple cells. To address this, ENACT offers three methods for assigning gene counts from shared bins to cells. In this study, the Weighted-by-Area method was applied. This approach assigns each shared bin *B*_*i*_ *∈ B*_shared_ to overlapping cells proportionally, based on the fraction of the bin’s area that intersects with each cell. The gene count assigned to cell *C*_*j*_ is computed as:

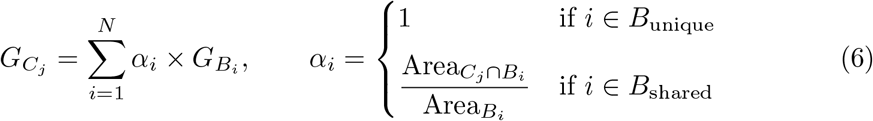

where 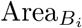 is the total area of bin *B*_*i*_ (*∼*4 *µ*m ), and Area_*C ∩B*_ is the intersection area between cell *C*_*j*_ and bin *B*_*i*_. For bins falling entirely within a single cell (*B*_unique_), *α*_*i*_ = 1, meaning their counts are assigned in full.

### 2.3 SpaGE Gene Imputation

Spatial transcriptomics platforms such as CosMx (*∼* 6,000 genes) and Xenium (*∼* 5,000 genes) capture only a subset of the transcriptome, leaving the majority of genes unmeasured. To recover whole-transcriptome expression in the spatial context, we applied SpaGE (Spatial Gene Enhancement) [30], a domain-adaptation-based imputation method that predicts the expression of spatially unmeasured genes by integrating the spatial data with a matched scRNA-seq reference dataset obtained from the same tissue. The SpaGE algorithm operates in two major steps: dataset alignment via the domain adaptation method PRECISE [38], followed by *k*-nearest-neighbour (kNN) gene expression prediction.

#### Step 1 - Dataset Alignment via PRECISE

Let *R ∈* ℝ ^*n×g*^ be the reference matrix for scRNA-sequence (*n* cells, *g* genes) and *Q ∈* ℝ ^*m×h*^ be the spatial query matrix (*m* cells, *h* genes). Using the set of shared genes *p* = *g ∩ h*, PRECISE independently applies Principal Component Analysis (PCA) to each data set, producing two sets of principal components *PC*_*r*_ and *PC*_*q*_. These are aligned by Singular Value Decomposition (SVD) of their cross-covariance matrix:

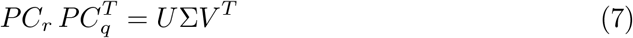

The resulting orthogonal transformations *U* and *V* are used to compute the so-called *Principal Vectors* (PVs):

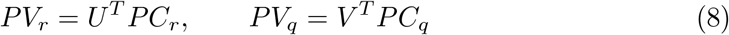

The PVs are sorted in decreasing order of cosine similarity, and only the informative PVs with cosine similarity *>* 0.3 are retained (effective number *d*^*′*^), discarding noisy or dataset-specific components. Both datasets are then projected onto *PV*_*r*_ to obtain a common aligned latent space:

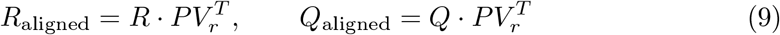

#### Step 2 - kNN Gene Expression Prediction

In the aligned latent space, for each spatial cell *i*, we identify its *k* = 50 nearest neighbors from the scRNA-seq reference using cosine distance. Only neighbors with positive cosine similarity (cosine distance *<* 1) are retained. The predicted expression *ŷ*_*il*_ of spatially unmeasured gene *l* for spatial cell *i* is then computed as:

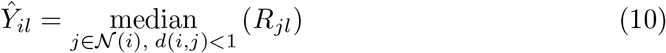

where *N* (*i*) denotes the set of nearest-neighbor scRNA-seq cells of spatial cell *i*, and *R*_*jl*_ is the expression of gene *l* in reference cell *j*. Note that we use the median rather than the weighted average proposed in the original publication [30]. This modification promotes sparsity in the imputed matrix by producing more exact zero values, which is biologically appropriate given the high dropout rates expected in single-cell data, and reduces the number of spurious non-zero (NNZ) entries that would otherwise arise from averaging across neighbors.

### 2.4 Evaluation

Tumour-label precision was evaluated using the author-provided cell-type annotations. Predicted tumour cells were considered supported when assigned to an epithelial compartment. Precision was computed as

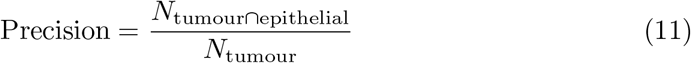

where *N*_tumour_ denotes the number of cells predicted as tumour and *N*_tumour*∩*epithelial_ denotes predicted tumour cells annotated as epithelial.

### 2.5 Implementation Details

The SpaceMACE pipeline is implemented entirely in R. The core MACE framework, including scATOMIC and SCEVAN, together with the spatial refinement model, are available as part of the SpaceMACE R package. This consolidation means that once the preprocessing steps have been completed, the full classification and refinement workflow can be run from within a single environment.

SpaGE and ENACT are not part of the SpaceMACE package and must be run as separate preprocessing steps before the pipeline is invoked. SpaGE is available as a Python package and was applied to imaging-based datasets prior to MACE classification to predict transcriptome-wide expression profiles of the targeted gene panels. ENACT is a standalone Python library designed for Visium HD data and was used to generate segmentation-informed cell-like units from the raw 2 *µ*m Visium HD input prior to MACE classification. In both cases, the output of the preprocessing step was passed directly as input to the SpaceMACE pipeline.

The fastcluster modification to SCEVAN replaces the internal call to stats::hclust with fastcluster::hclust, a drop-in replacement that requires no changes to the surrounding logic or decision thresholds and is incorporated directly into the SpaceMACE package.

## 3 Results

### 3.1 SpaceMACE Overview

To enable tumour/normal annotation of spatially resolved transcriptomics data, we developed SpaceMACE (Fig. 1). Accurate delineation of tumour and normal cells in spatial transcriptomics is challenging because existing tools such as MACE were designed for dissociated single-cell RNA sequencing data and cannot be directly applied to spatial platforms, which differ substantially in resolution, gene coverage, and data structure. To overcome these limitations, we extended MACE to four spatial transcriptomics platforms spanning two major modalities: sequencing-based ST (Stereo-seq and Visium HD) and imaging-based ST (CosMx and Xenium).

**Fig. 1.**
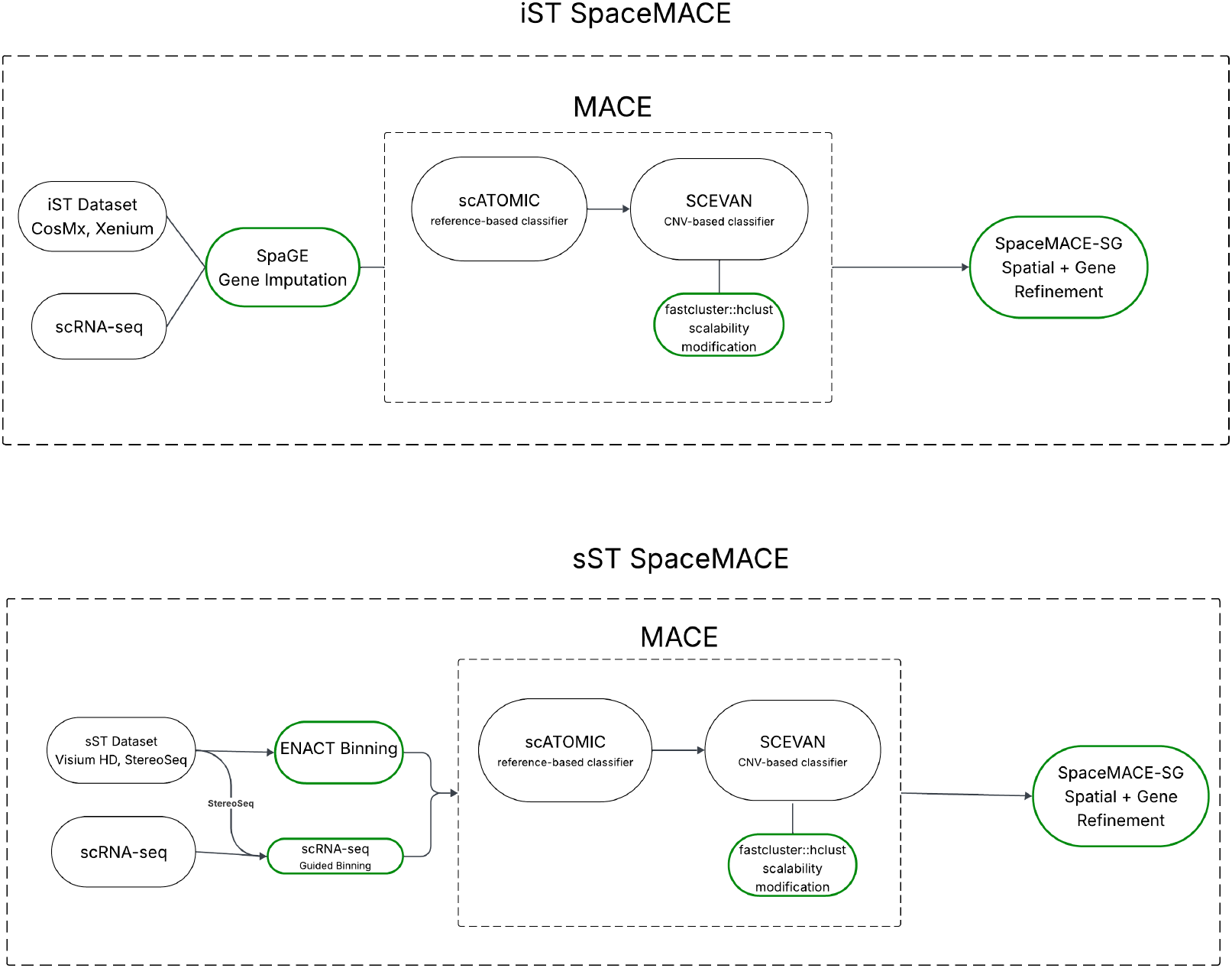
Overview of the SpaceMACE pipeline for imaging-based (top) and sequencing-based (bottom) spatial transcriptomics. Green outlines indicate novel contributions to the original MACE framework: SpaGE gene imputation for iST platforms, ENACT binning and scRNA-seq-guided binning for sST platforms, the fastcluster scalability modification to SCEVAN, and the SpaceMACE spatial refinement step.

For sequencing-based platforms, raw pixel or spot measurements are too sparse for reliable classification. We therefore evaluated two binning strategies to aggregate neighboring measurements into biologically meaningful units before applying MACE. The first strategy uses matched scRNA-seq data from the same tissue to select a bin size whose UMI distribution aligns with the single-cell reference. The second strategy uses ENACT, a segmentation-informed binning method that assigns Visium HD counts to putative cells using the tissue image, removing the requirement for a matched scRNA-seq reference. To support the larger cell counts encountered in spatial datasets, MACE’s internal clustering step was modified by replacing the standard hierarchical clustering routine in SCEVAN with fastcluster, a memory-efficient drop-in replacement that removes a hard cell-count ceiling and allows the full dataset to be processed in a single pass.

For imaging-based platforms, CosMx and Xenium provide near single-cell resolution but measure only targeted gene panels of a few thousand genes, which is insufficient for the copy-number inference component of MACE. To address this, we applied SpaGE, a domain-adaptation-based imputation method that predicts whole-transcriptome expression in spatial cells using matched scRNA-seq as a reference, expanding the targeted panels to a genome-wide representation compatible with MACE.

Finally, to leverage the spatial organization of cells within the tissue, we developed two spatial refinement models applied as a post-processing step on the MACE consensus output. SpaceMACE-S reassigns labels based on the majority vote among spatially proximal neighbors, while SpaceMACE weights neighbors by a combination of physical distance and gene expression similarity, making the refinement sensitive to both positional and transcriptional context. Together, these components form the SpaceMACE pipeline, which was evaluated across three cancer types — colorectal adenocarcinoma (COAD), hepatocellular carcinoma (HCC), and ovarian cancer (OV) — using datasets from the SPATCH resource with matched scRNA-seq and author-provided cell-type annotations.

### 3.2 MACE enables tumour-cell annotation in sequencing-based spatial transcriptomics

Sequencing-based spatial transcriptomics platforms measure broad, near-whole-transcriptome expression profiles and are therefore compatible with the gene-coverage requirements of MACE. However, these technologies often generate measurements at subcellular or near-cellular resolution, where individual pixels may contain too few transcripts for reliable tumour/normal classification.

#### 3.2.1 scRNA-seq-guided bin-size selection for Stereo-seq

As an initial evaluation, MACE was applied directly to the COAD Stereo-seq dataset using the original spatial resolution provided by the authors. The dataset contained 361,260 spatial units. Direct application of MACE resulted in extensive filtering, with 221,603 of 361,260 units removed, corresponding to 61.34% of the dataset. Filtered units were concentrated in low-signal and off-tissue regions (Suppl. Fig. S1), indicating that expression sparsity was a major limitation when applying MACE directly to the original spatial representation.

We validated the MACE labelling framework using matched scRNA-seq data from the same donors. MACE achieved a precision of 98.7%, confirming that the framework performs well on single-cell datasets. These findings suggest that the extensive filtering observed in Stereo-seq was primarily caused by the low transcript content of individual spatial units rather than by a biological loss of tumour-associated signal.

To reduce sparsity, neighbouring Stereo-seq pixels were aggregated into larger spatial bins. The bin size was selected by comparing the distribution of total UMIs per spatial bin across candidate resolutions with the total UMI distribution of the matched scRNA-seq reference (Fig. 2a). The bin size producing the closest agreement with the scRNA-seq distribution was selected independently for each cohort. This resulted in bin sizes of 10 *µ*m for COAD, 12.5 *µ*m for OV, and 15 *µ*m for HCC.

**Fig. 2.**
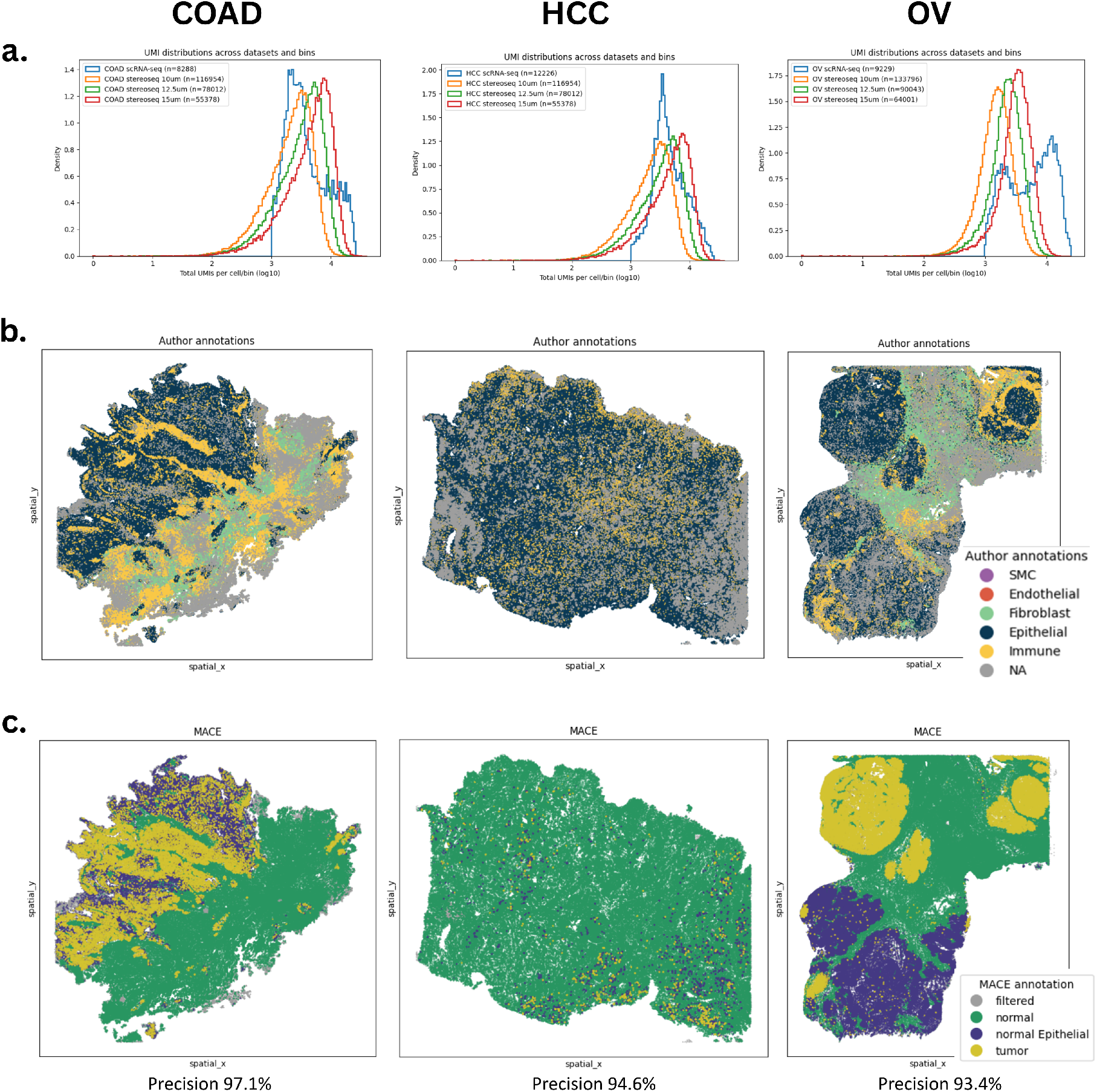
scRNA-seq-guided bin-size selection and MACE annotation of Stereo-seq datasets. **a**, UMI distributions across candidate spatial bin sizes compared with the matched scRNA-seq reference. Selected bin sizes were 10 *µ*m for COAD, 12.5 *µ*m for OV, and 15 *µ*m for HCC. **b**, Author-provided cell-type annotations harmonized into the major epithelial, immune, fibroblast, endothelial, smooth muscle cell, and unassigned compartments. **c**, MACE tumour/normal consensus annotations generated using the selected bin sizes. MACE achieved 97.1% precision in COAD, 94.6% in HCC, and 93.4% in OV.

The author-provided annotations were available at the original Stereo-seq pixel resolution and were collapsed into broad cellular compartments to support consistent interpretation across datasets (Fig. 2b). Epithelial cells and hepatocytes were grouped as epithelial, while fibroblasts, smooth muscle cells, and endothelial cells were retained as separate classes. Myeloid and lymphoid populations, including macrophages, mono-cytes, neutrophils, dendritic cells, mast cells, T cells, B cells, plasma cells, natural killer cells, and Kupffer cells, were grouped as immune.

MACE was then applied to the selected binned representation (Fig. 2c). To recover annotations at the native Stereo-seq resolution, each original pixel was assigned the consensus label of its parent bin. MACE achieved a precision of 97.1% in COAD, 94.6% in HCC, and 93.4% in OV. Despite the overall high precision, a subset of tumour-labelled pixels was associated with non-epithelial author annotations, indicating the presence of residual false-positive tumour calls outside the epithelial lineage.

#### 3.2.2 scRNA-seq-guided bin-size selection for Visium HD

An analogous analysis was performed for Visium HD. Candidate fixed-bin resolutions were compared with the matched scRNA-seq UMI distributions, resulting in selected bin sizes of 16 *µ*m for COAD, 16 *µ*m for OV, and 24 *µ*m for HCC (Fig. 3a). The author-provided cell-type annotations were harmonized into the same major cellular compartments used for Stereo-seq (Fig. 3b). MACE was then applied to the selected fixed-bin representations (Fig. 3c), achieving 87.7% precision in COAD and 98.7% precision in HCC. The OV dataset achieved approximately 90% precision. However, a dataset-specific issue caused the majority of spatial units to be filtered. Consequently, the OV result was excluded from the primary fixed-bin Visium HD evaluation and was not used to support the main conclusions.

**Fig. 3.**
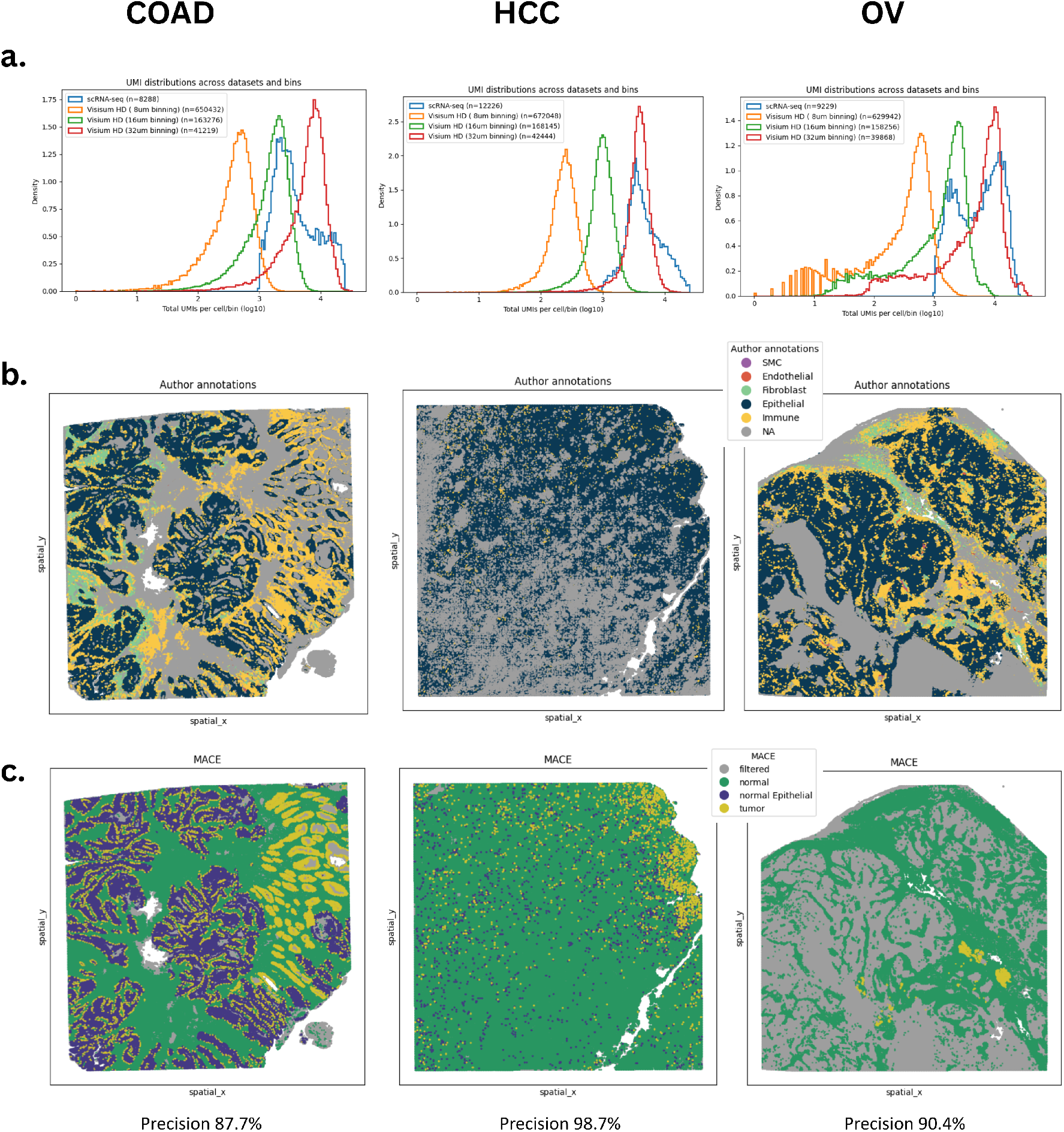
scRNA-seq-guided bin-size selection and MACE annotation of Visium HD datasets. **a**, UMI distributions at candidate 8, 16, and 32 *µ*m binning resolutions compared with the matched scRNA-seq reference. The selected bin size was the resolution most closely matching the scRNA-seq UMI distribution. **b**, Author-provided annotations harmonized into the major epithelial, immune, fibroblast, endothelial, smooth muscle cell, and unassigned compartments. **c**, MACE tumour/normal consensus annotations generated using the selected fixed-bin resolution. MACE achieved 87.7% precision in COAD and 98.7% in HCC. OV achieved approximately 90% precision but was excluded from the main fixed-bin evaluation because the majority of spatial units were filtered.

Although scRNA-seq-guided bin-size selection provides a principled way to choose a spatial resolution with transcript counts comparable to single-cell data, the strategy remains limited by the use of fixed square bins. Fixed binning may split the transcriptional profile of a single cell across neighbouring bins or merge expression from adjacent cells, particularly in regions with variable cell density or complex tissue architecture. In addition, matched scRNA-seq from the same sample or tissue section is not always available. We therefore evaluated ENACT as an alternative strategy for generating biologically informed spatial units without requiring matched scRNA-seq for bin-size selection.

#### 3.2.3 ENACT-derived spatial units improve Visium HD tumour-cell annotation

ENACT uses tissue images and Visium HD spatial coordinates to generate segmentation-informed cell-like units. Counts from the original 2 *µ*m Visium HD bins are reassigned to putative cells defined by morphological segmentation. Unlike fixed square binning, this strategy aims to produce spatial units that more closely reflect cellular boundaries and tissue organization.

We compared ENACT-derived units with scRNA-seq-guided fixed bins in terms of UMI distributions, matrix dimensions, and downstream MACE performance. In COAD, the ENACT-derived units showed a lower UMI peak than the matched scRNA-seq reference, reflecting the inclusion of smaller segmented objects. In contrast, 16 *µ*m fixed bins showed closer agreement with the scRNA-seq UMI distribution than 8 *µ*m bins (Suppl. Fig. S2).

These results highlight a trade-off between the two strategies. Fixed binning can be tuned to match a scRNA-seq UMI profile but requires an appropriate reference and does not respect cellular boundaries. ENACT does not require scRNA-seq-guided bin-size selection and instead produces morphology-informed spatial units, although these units may contain fewer transcripts than typical scRNA-seq cells.

The enhanced version of SCEVAN enabled MACE to process the complete ENACT-derived dataset in a single pass. This represented a substantial change from the previous implementation, in which large datasets had to be divided into smaller subsets because of the hierarchical clustering limit. Processing subsets independently prevented SCEVAN from accessing the global expression landscape and could alter the inferred copy-number structure.

The modified implementation used fastcluster and parallel distance computation, allowing SCEVAN to evaluate the complete dataset simultaneously. Consequently, the MACE output obtained with ENACT differed markedly from the output obtained with the earlier fixed-bin analysis (Suppl. Fig. S3). ENACT produced large, spatially coherent tumour regions that aligned with the epithelial tissue structure in the author-provided annotations.

Applying MACE to ENACT-derived Visium HD units yielded a precision of 97.8% in COAD and 99.1% in HCC (Fig. 4). This represented a substantial improvement over the fixed-bin COAD result of 87.7%, whereas the improvement in HCC was smaller because the fixed-bin result was already highly precise.

**Fig. 4.**
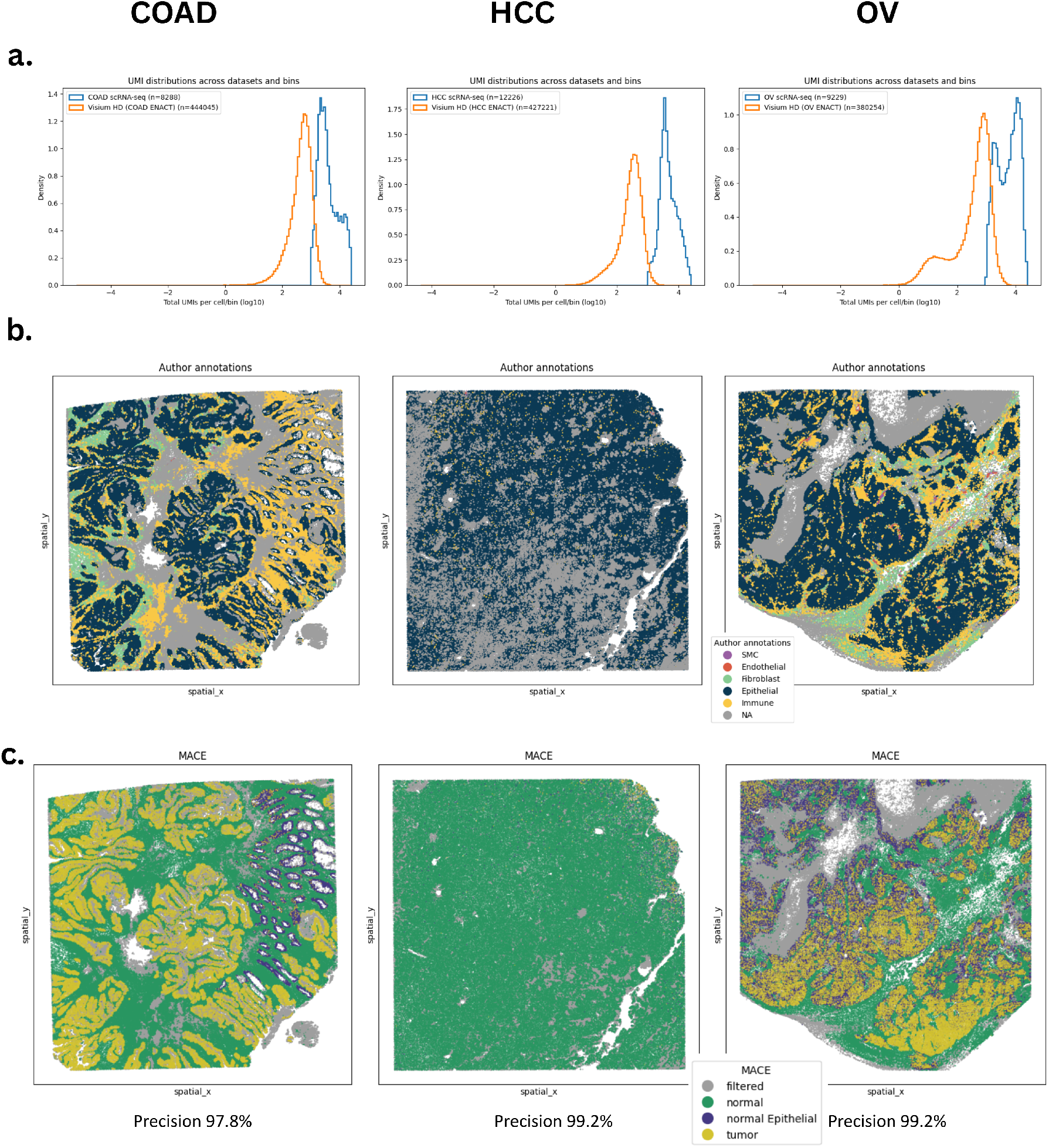
MACE annotation of ENACT-derived Visium HD spatial units. **a**, UMI distributions of ENACT-derived units compared with the matched scRNA-seq reference. **b**, Author-provided cell-type annotations harmonized into the major epithelial, immune, fibroblast, endothelial, smooth muscle cell, and unassigned compartments. **c**, MACE tumour/normal consensus annotations generated from ENACT-derived units. MACE achieved 97.8% precision in COAD and 99.1% in HCC.

Together, these results suggest that segmentation-informed spatial units provide a more suitable input for MACE than fixed square bins, particularly in morphologically heterogeneous tissues such as COAD.

### 3.3 SpaGE enables MACE analysis of imaging-based spatial transcriptomics

After evaluating sequencing-based platforms, we extended the analysis to the imaging-based CosMx and Xenium platforms. These technologies provide single-cell or near-single-cell resolution and therefore do not require spatial binning. However, both platforms measure targeted gene panels rather than the whole transcriptome, which is incompatible with the copy-number inference component of MACE, as it relies on broad chromosome-wide expression patterns. Direct application of MACE to the targeted panels was therefore not possible. To address this, SpaGE was used to predict genome-wide expression profiles using matched scRNA-seq data as a reference, providing the transcriptomic coverage required for MACE classification.

CosMx operates at single-cell resolution and is conceptually well aligned with the original MACE framework. Following SpaGE imputation, MACE achieved 99.9% precision across all three cohorts — COAD, HCC, and OV (Fig. 5). The same workflow was then applied to Xenium, which similarly measures targeted panels at near-single-cell resolution. MACE again achieved 99.9% precision in both COAD and HCC (Suppl. Fig. S4). The consistency of results across both platforms demonstrates that SpaGE imputation provides sufficient transcriptomic coverage for MACE while preserving highly accurate tumour-cell classification, and supports the applicability of the SpaGE–MACE workflow across imaging-based spatial transcriptomics platforms more broadly.

**Fig. 5.**
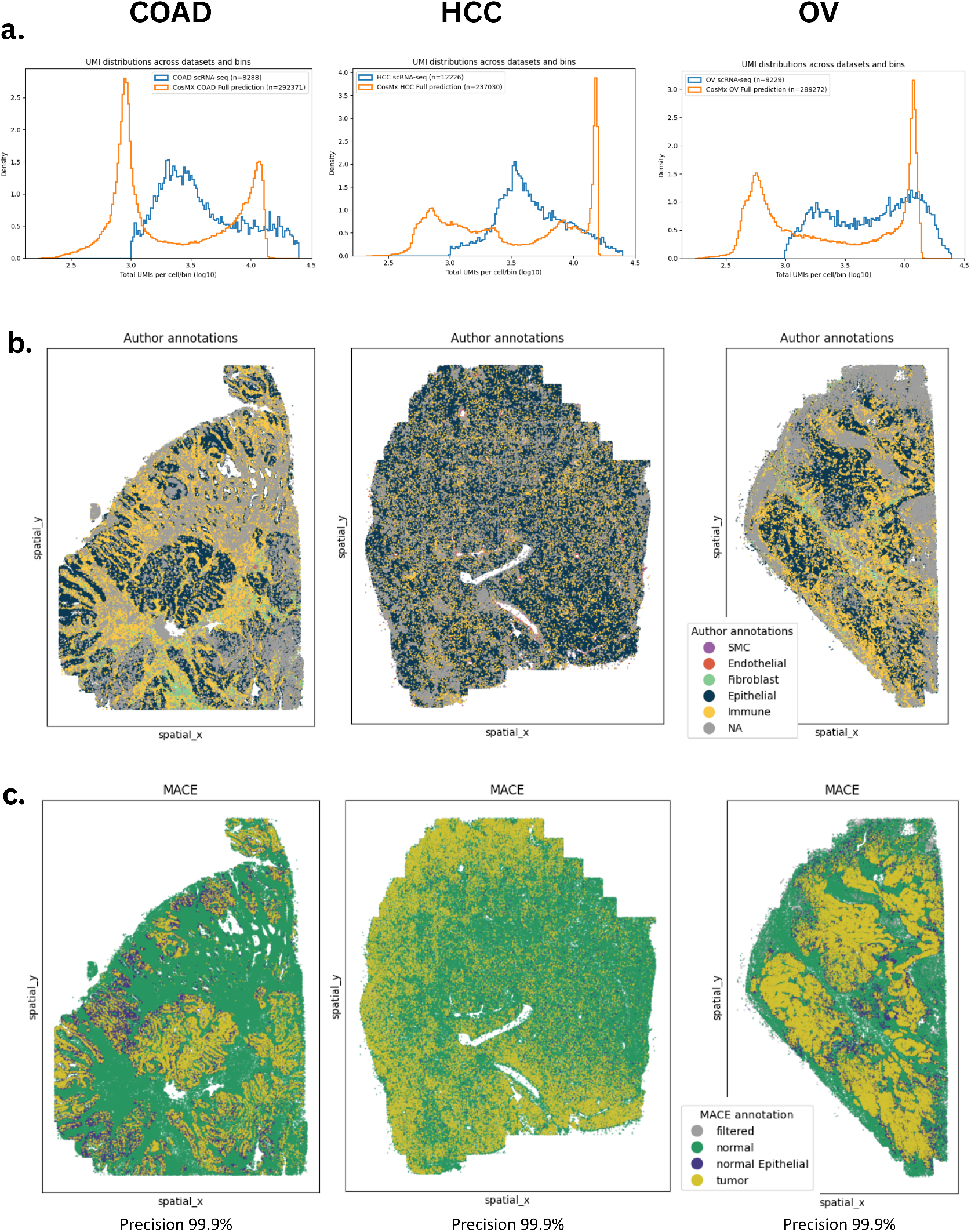
MACE tumour-cell annotation of SpaGE-imputed CosMx datasets. **a**, UMI distributions of CosMx cells following SpaGE imputation compared with the matched scRNA-seq reference. **b**, Author-provided annotations harmonized into the major epithelial, immune, fibroblast, endothelial, smooth muscle cell, and unassigned compartments. **c**, MACE tumour/normal consensus annotations generated from the SpaGE-imputed expression profiles. MACE achieved approximately 99.9% precision in COAD, HCC, and OV.

### 3.4 SpaceMACE incorporates spatial and transcriptomic information into MACE predictions

A limitation of the original MACE framework is that tumour/normal classification is based exclusively on gene-expression-derived evidence and does not explicitly account for the spatial organization of cells within the tissue. Tumour cells frequently form spatially coherent regions, while normal cells are often surrounded by cells with similar identities. We therefore developed SpaceMACE as a post-processing refinement of the MACE consensus labels. Two variants were evaluated. SpaceMACE-S uses spatial proximity alone, whereas SpaceMACE combines spatial proximity with gene-expression similarity to weight the contribution of neighbouring cells. Performance was evaluated using epithelial-support precision together with the number of retained tumour calls. Only datasets with complete MACE, SpaceMACE-S, and SpaceMACE outputs were included in the quantitative comparison. OV Xenium was excluded because the complete workflow was not available.

Table 2 summarizes tumour-cell precision and tumour counts across all evaluated platforms and datasets. In Visium HD with scRNA-seq-guided binning, the HCC slide was composed predominantly of normal cells and contained relatively few tumour calls. In this setting, precision alone is insufficient to assess performance: a high precision value can be achieved when only a small number of cells are called tumour, provided that these calls occur within the epithelial compartment. SpaceMACE reduced the HCC tumour count from 19,358 cells with MACE to 11,323 cells with SpaceMACE-S and 14,402 cells with SpaceMACE. Therefore, spatial refinement removed part of an already limited tumour-labelled population.

**Table 2.** Tumour-cell precision and number of tumour calls across MACE, SpaceMACE-S, and SpaceMACE.

|  |  | **MACE** |  | **SpaceMACE-S** |  | **SpaceMACE** |  |
|---|---|---|---|---|---|---|---|
| **Platform** | **Dataset** | **Precision** | **Tumour cells** | **Precision** | **Tumour cells** | **Precision** | **Tumour cells** |
| CosMx | COAD | **99.94%** | 26,640 | 99.38% | 20,884 | 99.89% | 29,303 |
|  | HCC | **99.98%** | 42,357 | 94.16% | 19,119 | 99.57% | 29,303 |
|  | OV | **99.93%** | 56,044 | 98.76% | 58,523 | 99.59% | 57,459 |
| Stereo-seq | COAD | 95.89% | 66,066 | 96.17% | 67,212 | **96.18%** | 66,458 |
|  | HCC | **94.58%** | 5,336 | 94.48% | 2,683 | 94.12% | 4,014 |
|  | OV | 93.41% | 73,963 | 93.68% | 74,412 | **93.69%** | 74,137 |
| Visium HD<br>ENACT | COAD | 97.80% | 64,319 | **98.03%** | 61,187 | 97.86% | 63,066 |
|  | HCC | **99.11%** | 568 | 97.67% | 129 | 98.01% | 151 |
|  | OV | **99.20%** | 42,637 | 98.90% | 33,048 | 99.04% | 37,466 |
| Visium HD<br>fixed bins | COAD | 87.74% | 54,318 | **89.34%** | 50,759 | 88.30% | 53,257 |
|  | HCC | 98.72% | 19,358 | **98.97%** | 11,323 | **98.97%** | 14,402 |
|  | OV | **99.20%** | 3,434 | 98.90% | 3,317 | 99.04% | 3,317 |
| Xenium | COAD | **99.95%** | 35,982 | 98.64% | 33,286 | 99.55% | 34,070 |
|  | HCC | **99.98%** | 68,805 | 97.34% | 38,333 | 99.92% | 61,297 |

*Note:* Precision represents the proportion of predicted tumour cells supported by the epithelial compartment in the harmonized reference annotation. Tumour counts should be interpreted together with precision, particularly in low-tumour-burden or extensively filtered datasets.

A similar caution applies to OV in the Visium HD fixed-bin analysis. Although precision remained high, extensive filtering left only a small number of tumour calls compared with the other OV datasets. The high precision should therefore be interpreted as strong epithelial support among the retained tumour calls rather than as evidence of high tumour recovery.

This behaviour is consistent with low-tumour-burden tissues. When tumour-labelled cells are sparse and surrounded by a large normal background, neighbourhood-based smoothing may reassign them to the dominant local normal label. This effect was stronger for SpaceMACE-S, which uses only spatial information, and weaker for SpaceMACE, which also incorporates gene-expression similarity.

A comparable pattern was observed in imaging-based HCC datasets. SpaceMACE-S substantially reduced the number of tumour calls in both CosMx and Xenium, whereas SpaceMACE retained a larger proportion of the original MACE tumour population. This indicates that spatial information alone may over-smooth tumour annotations, while the addition of gene-expression similarity produces a more conservative refinement.

Across sequencing-based datasets, SpaceMACE produced modest precision improvements in several cases. In Stereo-seq, SpaceMACE increased precision from 95.89% to 96.18% in COAD and from 93.41% to 93.69% in OV. SpaceMACE-S produced similar but slightly smaller improvements.

The largest improvement in fixed-bin Visium HD was observed in COAD, where precision increased from 87.74% with MACE to 89.34% with SpaceMACE-S and 88.30% with SpaceMACE. These results indicate that spatial refinement can correct a subset of locally inconsistent false-positive tumour calls in binned sequencing-based spatial data.

The effect of SpaceMACE was less consistent in imaging-based datasets. CosMx and Xenium already achieved near-perfect MACE precision before spatial refinement.

Consequently, additional spatial smoothing provided little benefit and, in several cases, reduced precision. The strongest decrease occurred in CosMx HCC, where precision fell from 99.98% with MACE to 94.16% with SpaceMACE-S. SpaceMACE was more conservative and retained 99.57% precision, but still did not improve over the original MACE output. These results suggest that SpaceMACE is most beneficial when the initial MACE output contains local spatial noise. When the original single-cell classification is already highly accurate, additional spatial smoothing may reduce performance.

#### 3.4.1 Label transitions induced by SpaceMACE

To investigate how SpaceMACE changed the original MACE output, label transitions were quantified between the original and refined annotations.

Transitions were grouped according to their expected effect on the epithelial-support precision metric. Precision-increasing transitions were classified as correct, precision-decreasing transitions as incorrect, and ambiguous transitions as neutral. Neutral transitions were not directly scored as correct or incorrect under the epithelial-support metric, although they could affect the total number of retained tumour calls.

Visium HD ENACT COAD was selected as a representative dataset for detailed inspection of label-flow patterns. Sankey diagrams show the relationship between author-provided epithelial or non-epithelial annotations, the original MACE labels, and the labels produced by SpaceMACE-S and SpaceMACE (Fig. 6). In the ENACT COAD dataset, both SpaceMACE variants produced more precision-increasing than precision-decreasing transitions. The dominant neutral transition consisted of epithelial cells moving from tumour to normal epithelial. This transition does not represent correction of a false-positive tumour call under the epithelial-support metric because the cells remain within the epithelial lineage. However, it reduces the total number of tumour calls.

**Fig. 6.**
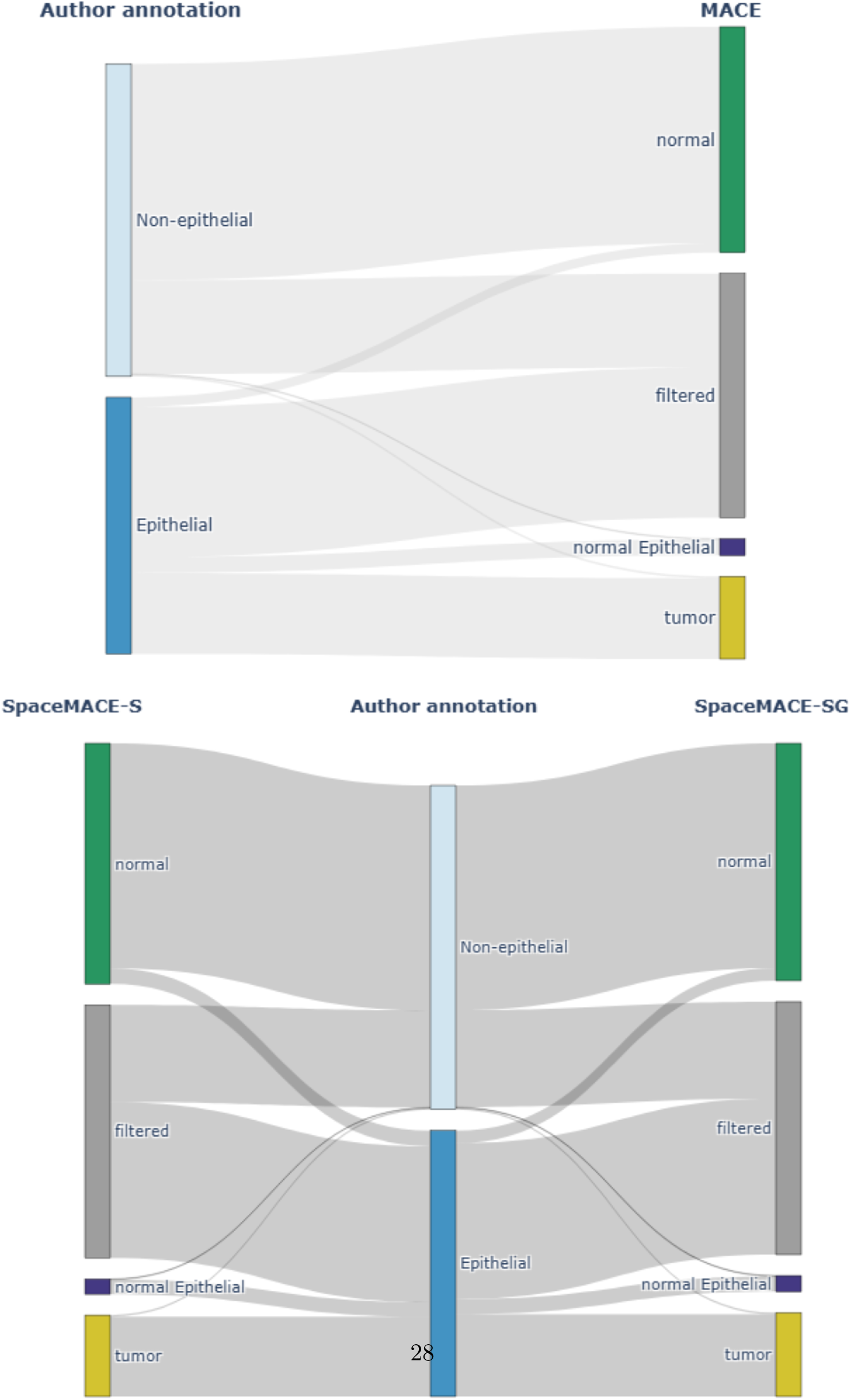
Label-flow analysis for Visium HD ENACT COAD. Sankey diagrams show the relationship between author-provided epithelial or non-epithelial annotations, the original MACE labels, and the SpaceMACE-refined labels. The upper panel shows transitions from the author annotation to the original MACE output. The lower panels compare SpaceMACE-S, left, and SpaceMACE, right, and show how each refinement strategy changes the original MACE labels.

SpaceMACE-S induced more relabelling events than SpaceMACE, whereas SpaceMACE changed fewer cells and retained a larger proportion of the original tumour population. Transition patterns were also summarized at the dataset level for Visium HD ENACT COAD and HCC (Fig. 7). HCC showed a different transition pattern because the original MACE output contained very few tumour-labelled cells relative to the normal population. As a result, many tumour-labelled cells were surrounded by normal neighbours and were reassigned to normal or normal epithelial, particularly by SpaceMACE-S. This explains the marked reduction in tumour-cell count and the absence of a precision improvement in HCC.

**Fig. 7.**
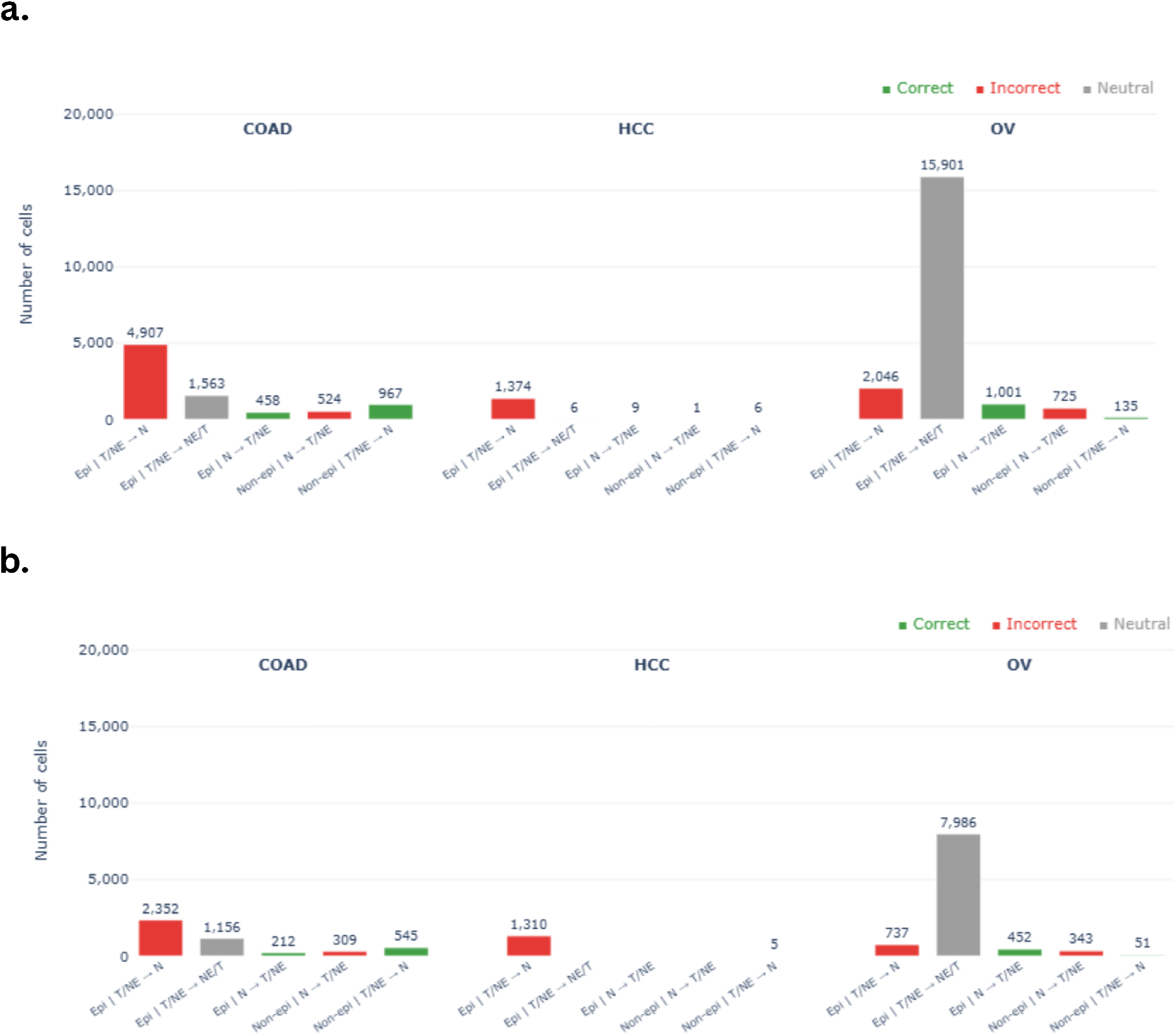
Classification of SpaceMACE-induced label transitions in Visium HD ENACT datasets. Bar charts show the number of cells undergoing label transitions in COAD, HCC and OV for **a** SpaceMACE-S and **b** SpaceMACE. Transitions are stratified by epithelial or non-epithelial reference annotation and by their expected effect on epithelial-support precision. Precision-increasing transitions are shown as correct, precision-decreasing transitions as incorrect, and ambiguous epithelial transitions as neutral. T, tumour; NE, normal epithelial; N, normal.

Detailed Sankey diagrams and transition summaries for the remaining platforms and datasets are provided in the Supplementary Information (Suppl. Fig. S5-S20).

## 4 Discussion

In this work, we extended MACE, a tool originally designed for single-cell RNA sequencing datasets, to a broader pipeline capable of annotating cells within the tumour micro-environment on both sequencing-based and imaging-based spatial transcriptomics platforms. The core contribution is SpaceMACE, a pipeline that integrates spatial context into the tumour/normal classification task, going beyond what the original MACE framework could offer by incorporating the physical location of cells within the tissue. SpaceMACE achieves platform compatibility through two key preprocessing strategies: SpaGE-based expression imputation for imaging-based platforms such as CosMx and Xenium, which lack whole-transcriptome coverage, and ENACT-based segmentation-informed binning for sequencing-based platforms such as Visium HD, which require aggregation of capture areas into biologically meaningful spatial units. Together, these adaptations allow MACE to be applied consistently across platforms that differ substantially in resolution, gene coverage, and data structure. The spatial refinement itself is implemented in two complementary models: SpaceMACE-S, which uses spatial neighbourhood information alone to enforce local label consistency, and SpaceMACE, which combines spatial proximity with gene expression similarity for a richer and more conservative refinement. These results provide a stable foundation for the discovery of tumour-specific targets in spatially resolved data. However, understanding when and where these contributions are most effective requires a closer look at the results across platforms and at the practical constraints of the current implementation. Fig. 1 provides an overview of the two SpaceMACE pipelines developed in this work, highlighting the novel contributions introduced relative to the original MACE framework.

Based on the results presented here, SpaceMACE is the recommended variant across both types of platform. For sequencing-based spatial transcriptomics, both SpaceMACE-S and SpaceMACE produced consistent precision gains over MACE, with SpaceMACE achieving the highest precision while making fewer total label changes. For imaging-based platforms, neither variant improved over the already nearperfect MACE output, and SpaceMACE-S in particular reduced precision in several cases by over-smoothing spatially isolated tumour calls. SpaceMACE was more conservative and better preserved the original MACE output in these settings. The spatial refinement step is therefore most valuable for sequencing-based data, where binning introduces local label noise that neighbourhood-based smoothing can correct. For imaging-based data, the primary contribution of SpaceMACE is not the refinement step itself but rather the enabling infrastructure: SpaGE imputation and fastcluster clustering now allow the full SpaceMACE pipeline to be applied consistently across all four platforms evaluated here without major platform-specific modifications.

All results in this work are derived from SPATCH datasets, and the number of datasets available per platform and type of cancer remains limited. Although the observed precision values are encouraging, they should be interpreted with caution given this constraint. STAGE [39], which profiles tumours on additional spatial transcriptomics platforms with matched author annotations, represents a natural extension dataset; however, its full release was pending at the time of writing. The addition of STAGE data, which include Visium HD slides with author annotations per-cell, would already significantly expand the evaluation and allow a more robust assessment of whether the trends observed here generalise across cohorts and tissue types. Beyond dataset coverage, the current implementation also carries several technical limitations that constrain its applicability.

The most immediate constraint is computational. Processing datasets exceeding 300,000 cells requires runtimes on the order of three days and peak memory usage of approximately 500 GB of RAM, with some cases exceeding this. Without access to a high-memory computing cluster, it is not possible to run the entire pipeline on a single data set in one pass, representing a significant barrier to accessibility and reproducibility outside of well-resourced environments. A second limitation concerns platform coverage: equivalent ENACT-based binning could not be completed for Stereo-seq within the scope of this work. There are two pathways for future exploration. The first is to adapt the Stereo-seq dataset to the input format expected by ENACT, which appears to be more tractable than modifying the ENACT library itself. The second and potentially more natural alternative is CellBin2, the upgraded version of Cell-Bin [40] developed by STOmics as part of their Stereo-seq Analysis Workflow (SAW) processing pipeline, which is designed natively for Stereo-seq data and could serve as a direct equivalent to ENACT for segmentation-informed binning on this platform.

Addressing these limitations defines the primary roadmap for future development. The most pressing need is to reduce the runtime and memory demands of SCEVAN for large datasets, either by identifying more scalable alternative libraries or by developing an internal tool optimised for high-cell-count spatial transcriptomics data. Resolving the Stereo-seq binning gap, whether through ENACT adaptation or CellBin2 integration, would complete platform coverage and allow SpaceMACE to be benchmarked consistently across all four platforms evaluated here.

In parallel, residual filtration in ENACT-binned Visium HD data and, prospectively, in a future Stereo-seq segmentation pipeline remains an open problem. ALRA [41], a zero-preserving imputation method, represents a possible strategy to recover low-signal cells before MACE classification. However, its adoption would require careful evaluation: the computational cost is substantial, and introducing an additional imputation layer before SCEVAN adds uncertainty to the expression profiles that the CNV inference relies on, which could compromise classification quality. Therefore, any future integration of ALRA into the pipeline should be preceded by a systematic assessment of whether the recovered cells actually improve tumour / normal classification or introduce noise that outweighs the benefit of reduced filtration.

Finally, although the current SpaceMACE pipeline is available as R package, parts of the preprocessing tools such as ENACT or SpaGE are Python based, which adds friction to deployment of a single pipeline and limits parallelisation options. As spatial transcriptomics datasets continue to grow in scale and complexity, consolidating the pipeline into a single language would improve maintainability, scalability, and the ability to leverage modern deep learning frameworks for future refinements of the spatial classification models.

## Supplementary information

Supplementary information accompanies this article and contains additional figures and dataset-level results.

## Declarations

### Data availability

All datasets analysed in this study are publicly available and were obtained from the SPATCH database https://spatch.pku-genomics.org/

### Code availability

The SpaceMACE source code is currently available at: https://gitlab.com/genmab-public/spaceMACE

