## Supplementary material for "SpaceMACE: Spatially-Aware Tumour Cell Annotation for Spatial Transcriptomics": Suppl. Fig. S

[1] Discovery Data Science, Genmab B.V., Utrecht, The Netherlands .

**Abstract**

This document provides the supplementary material for spaceMACE. An extension of the MACE tumour-cell annotation framework for sequencing-based and imaging-based spatial transcriptomics. SpaceMACE first combines reference-based cell-type classification with copy-number-variation-based tumour classification. The resulting consensus annotations are subsequently refined using spatial proximity, gene-expression similarity, or a combination of both.
We evaluated SpaceMACE across multiple spatial transcriptomics platforms, including Stereo-seq, Visium HD, CosMx and Xenium, and across colorectal, hepatocellular and ovarian cancer datasets. SpaceMACE retained the high precision of the original MACE framework while improving the spatial coherence of tumour-cell annotations. Its performance depended on spatial resolution, tumour burden, transcriptomic coverage and the preprocessing strategy used to define spatial observations.
Together, these results demonstrate that SpaceMACE provides a scalable and platform-independent framework for tumour-cell annotation in spatial transcriptomics and supports the spatial investigation of tumour-specific molecular targets.

**Keywords:** spatial transcriptomics, tumour-cell annotation, copy-number variation, single-cell RNA sequencing, spatial neighbourhood, target discovery

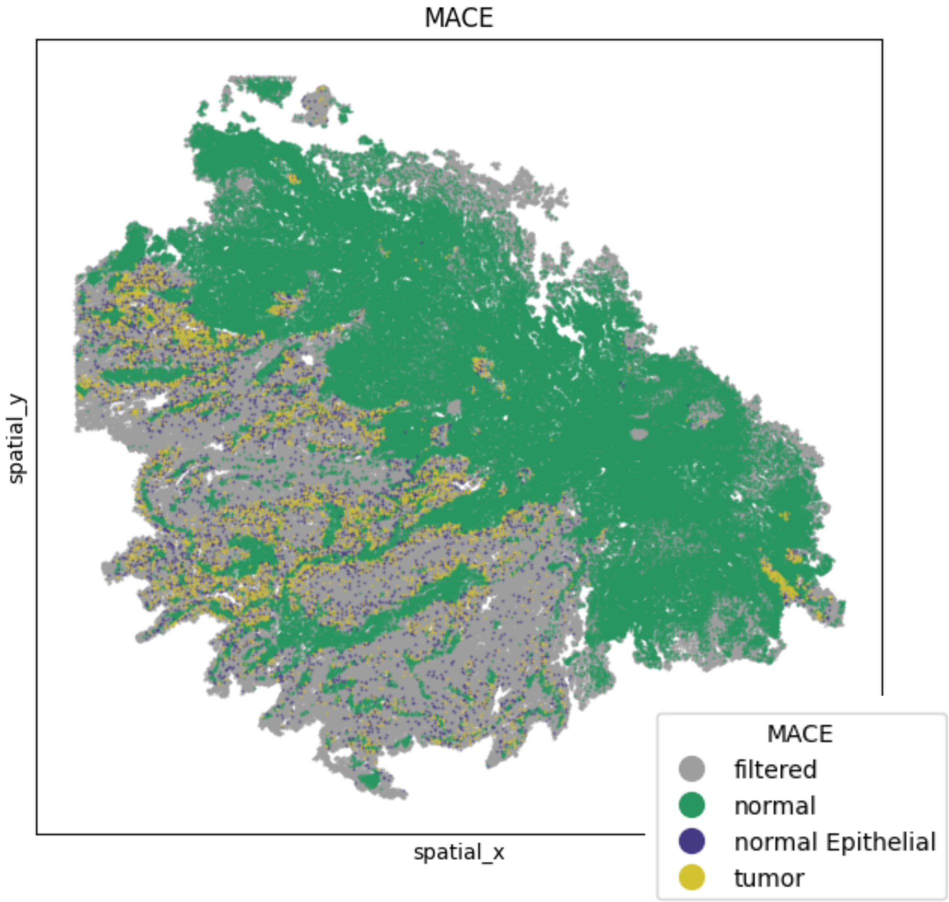


**Suppl. Fig. S1 Direct application of MACE to the original COAD Stereo-seq representation.** MACE tumour/normal annotation of COAD Stereo-seq data before additional spatial binning. A large proportion of the original spatial units was filtered because of insufficient transcript counts. Filtered units were concentrated primarily in low-signal and off-tissue regions.

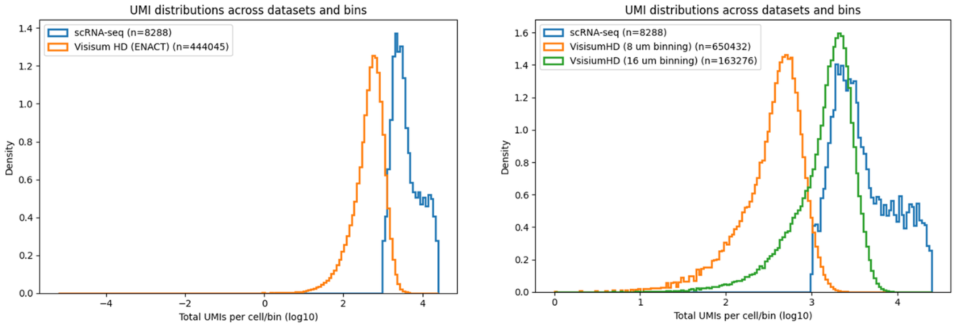


**Suppl. Fig. S2 Comparison of ENACT-derived spatial units and scRNA-seq-guided fixed bins in Visium HD COAD.** Left, UMI distribution of ENACT-derived units compared with the matched scRNA-seq reference. The ENACT distribution peaks at lower UMI counts, reflecting the inclusion of smaller segmented units. Right, UMI distributions obtained using fixed 8 and 16 $\mu$m bins. The 16 $\mu$m representation more closely matches the scRNA-seq UMI distribution than the 8 $\mu$m representation. Fixed binning can be tuned to match scRNA-seq transcript counts but requires a matched reference, whereas ENACT produces morphology-informed spatial units without scRNA-seq-guided bin-size selection.

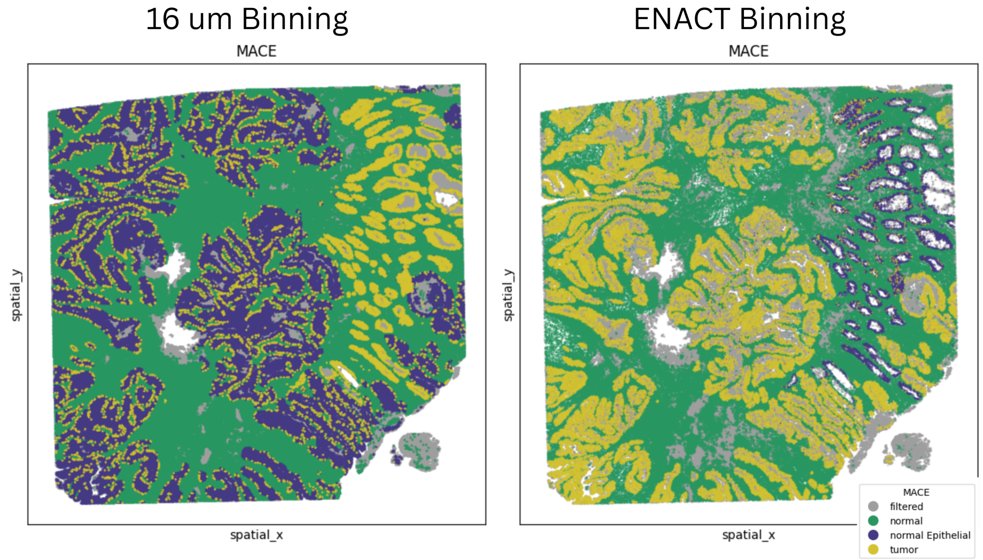


**Suppl. Fig. S3 Comparison of fixed-bin and ENACT-derived MACE annotations in Visium HD COAD.** MACE tumour/normal consensus annotations generated using scRNA-seq-guided 16 $\mu$m fixed bins, left, and ENACT-derived spatial units, right. The ENACT representation produces large, spatially coherent tumour regions that align with the epithelial tissue structure in the author-provided annotations. The enhanced SCEVAN implementation allowed the complete ENACT dataset to be processed simultaneously, providing access to the global expression landscape for copy-number inference.

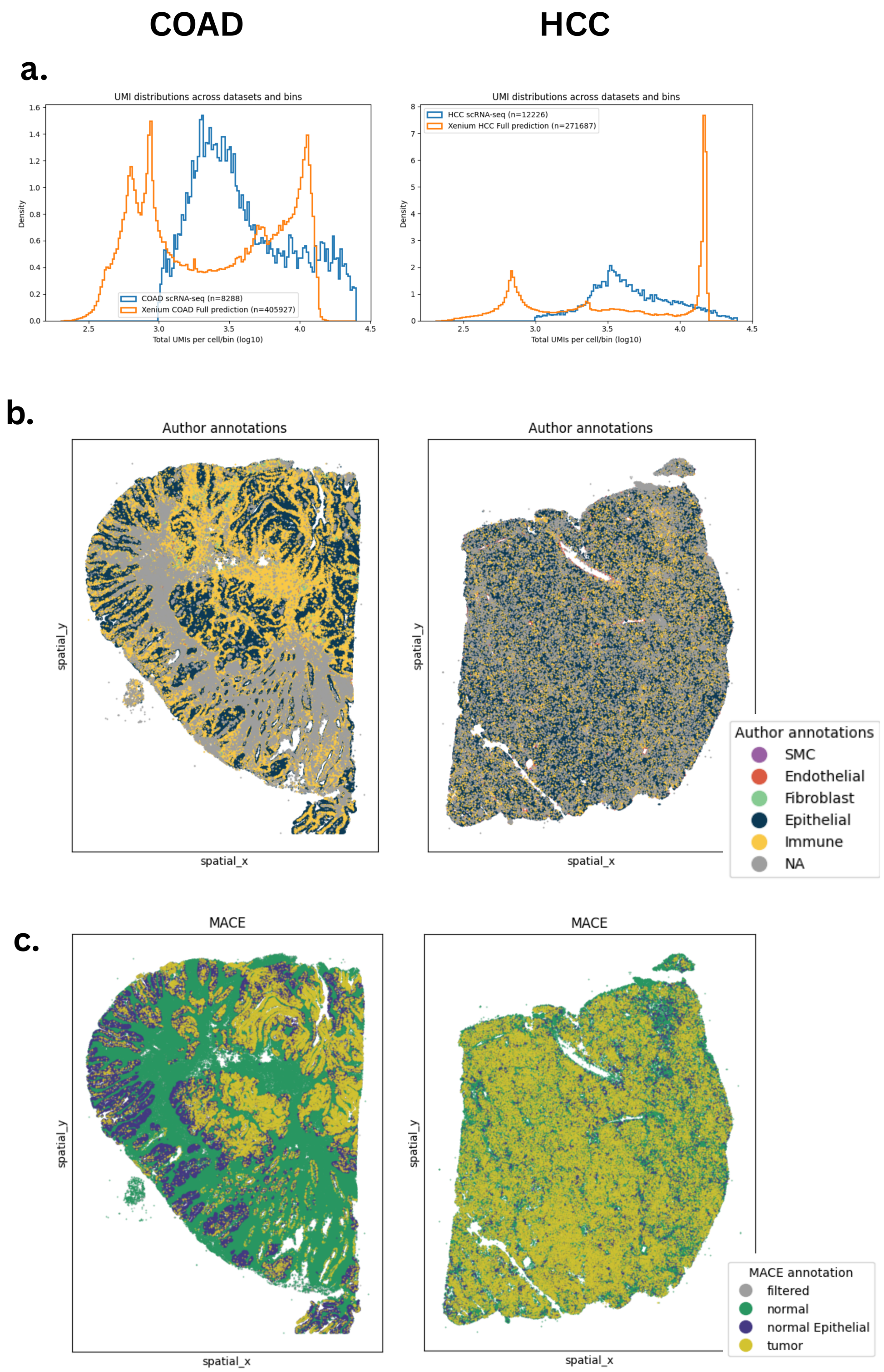




**Suppl. Fig. S4 MACE tumour-cell annotation of SpaGE-imputed Xenium datasets.** **a**, UMI distributions of Xenium cells following SpaGE imputation compared with the matched scRNA-seq reference. **b**, Author-provided annotations harmonized into the major epithelial, immune, fibroblast, endothelial, smooth muscle cell, and unassigned compartments. **c**, MACE tumour/normal consensus annotations generated from the SpaGE-imputed expression profiles. MACE achieved approximately 99.9% precision in COAD and HCC.

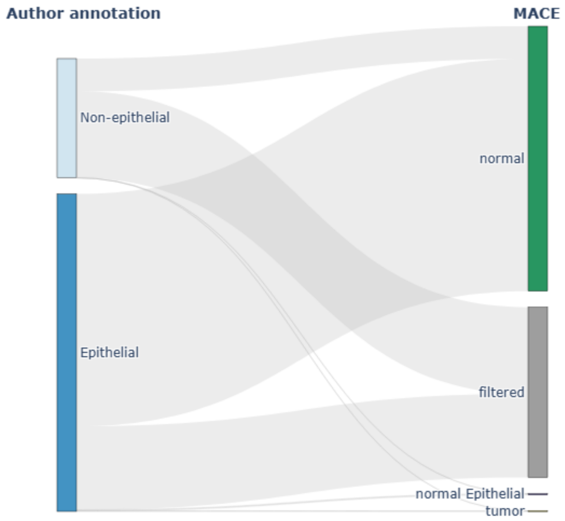


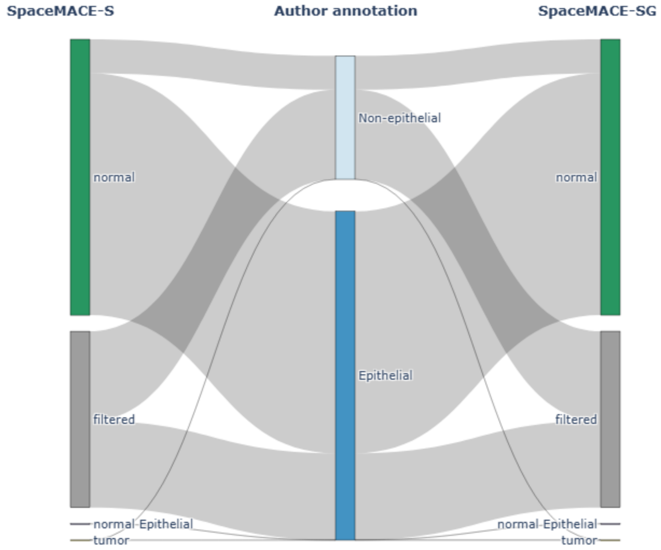


**Suppl. Fig. S5 Detailed label-flow analysis for VisiumHD ENACT HCC.** Sankey diagrams show the relationship between author-provided epithelial/non-epithelial annotations, the original MACE labels, and SpaceMACE-refined labels. *(Top)* Flow from author annotation to MACE output. *(Bottom)* Comparison of SpaceMACE-S (left) and SpaceMACE (right) outputs relative to the author annotation. The dominant transition is epithelial cells reassigned from tumor to normal, consistent with the low-tumor-burden pattern described for HCC.

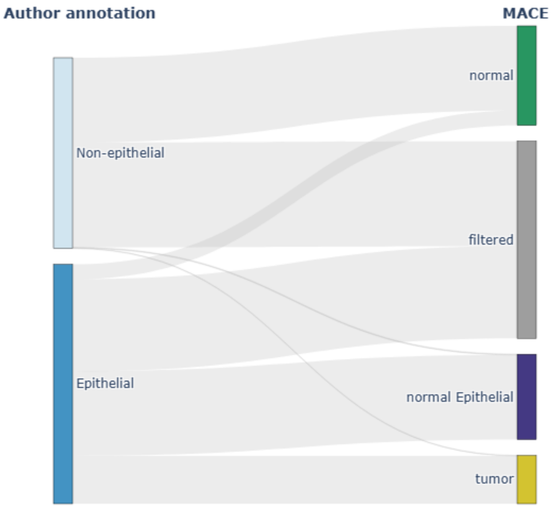


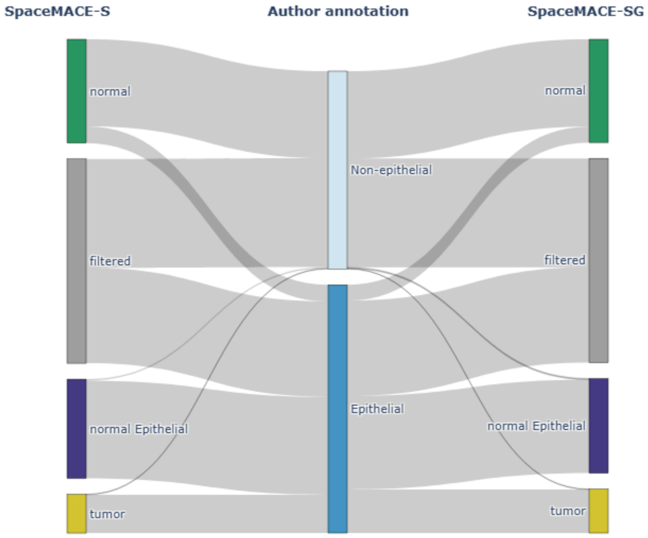


**Suppl. Fig. S6 Detailed label-flow analysis for VisiumHD ENACT OV.** Sankey diagrams show the relationship between author-provided epithelial/non-epithelial annotations, the original MACE labels, and SpaceMACE-refined labels. *(Top)* Flow from author annotation to MACE output. *(Bottom)* Comparison of SpaceMACE-S (left) and SpaceMACE (right) outputs relative to the author annotation. The dominant transition is epithelial cells reassigned from tumor to normal, consistent with the low-tumor-burden pattern described for HCC.

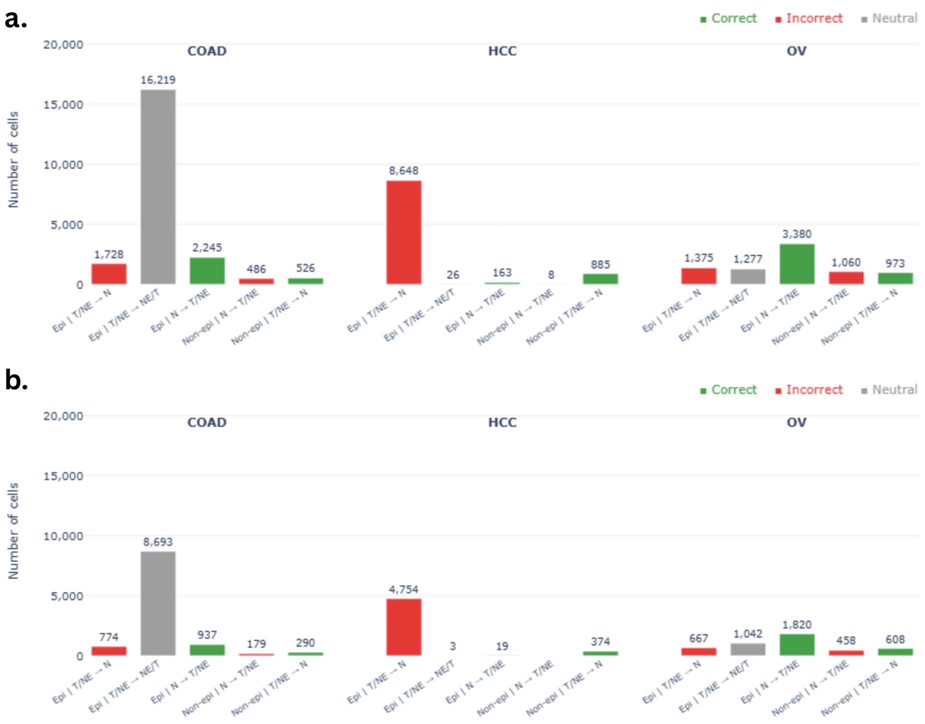


**Suppl. Fig. S7  Cell transition classification across cancer types for SpaceMACE-S and SpaceMACE on Stereo-seq data.** Bar charts showing the number of cells undergoing spatial domain transitions in colorectal adenocarcinoma (COAD), hepatocellular carcinoma (HCC), and ovarian cancer (OV), stratified by cell type (epithelial vs. non-epithelial) and transition direction. Transitions are colored by classification outcome: red (Incorrect) indicates cells reassigned to an incorrect spatial domain; green (Correct) indicates cells correctly reassigned; gray (Neutral) indicates ambiguous transitions within the epithelial compartment. T, tumor; NE, normal epithelial; N, normal. **(a)** SpaceMACE-S. **(b)** SpaceMACE.

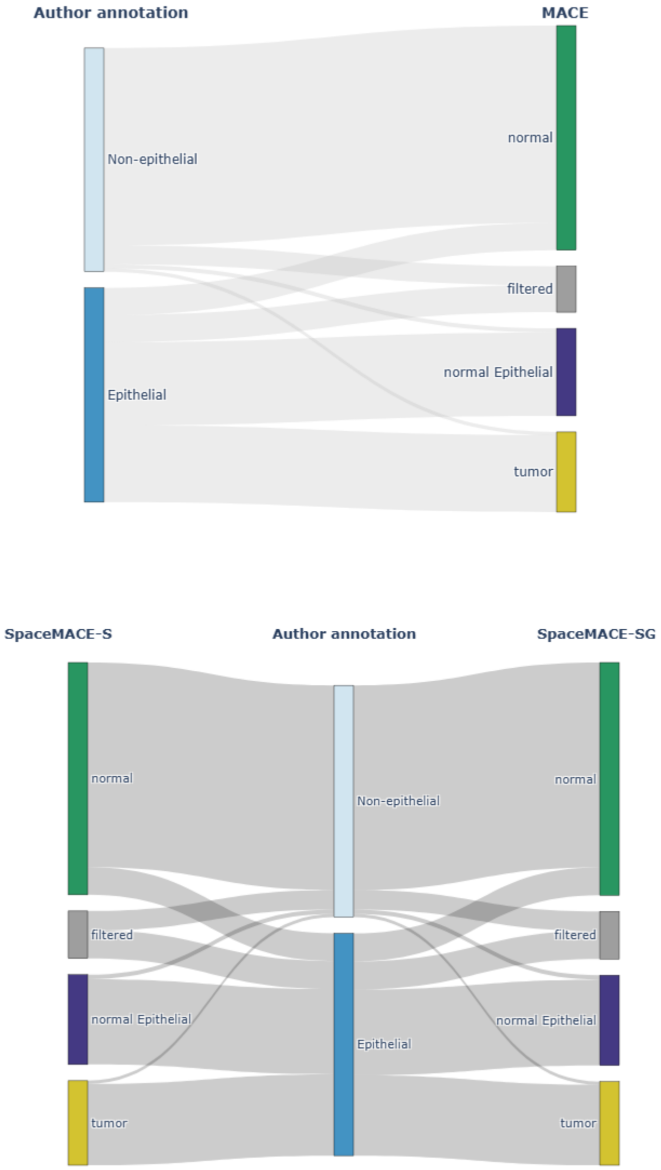


**Suppl. Fig. S8 Detailed label-flow analysis for Stereo-seq COAD.** Sankey diagrams show the relationship between author-provided epithelial/non-epithelial annotations, the original MACE labels, and SpaceMACE-refined labels. *(Top)* Flow from author annotation to MACE output. *(Bottom)* Comparison of SpaceMACE-S (left) and SpaceMACE (right) outputs relative to the author annotation. Both variants show a predominance of precision-increasing transitions, with SpaceMACE producing fewer total relabeling events.

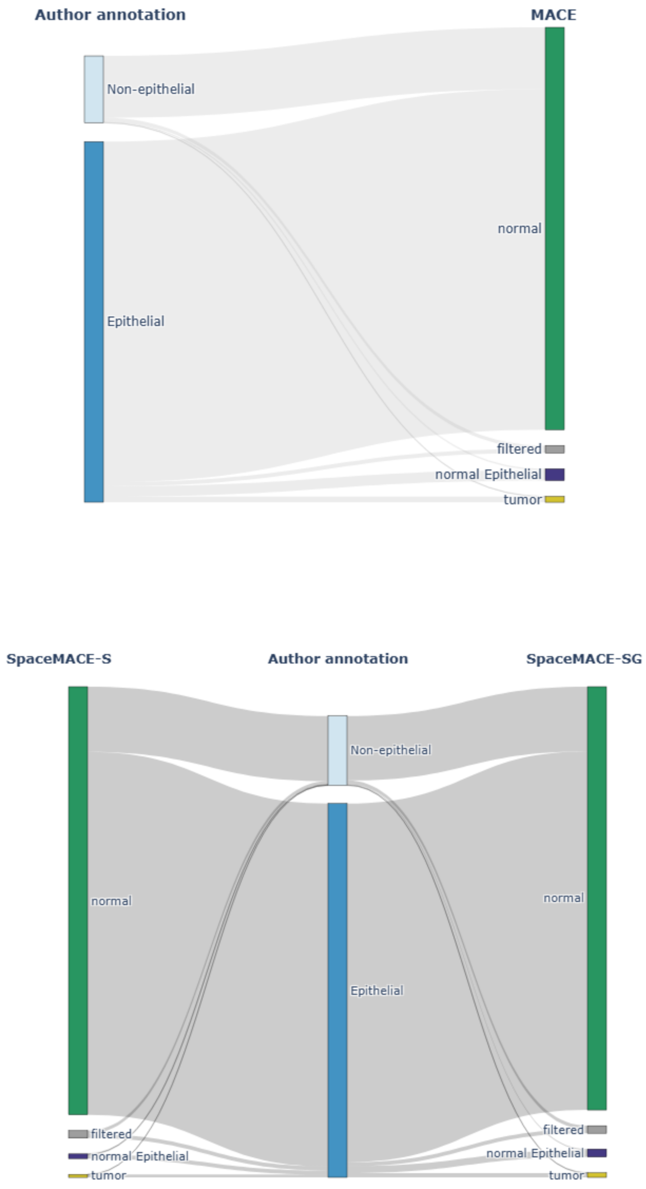


**Suppl. Fig. S9 Detailed label-flow analysis for Stereo-seq HCC.** Sankey diagrams show the relationship between author-provided epithelial/non-epithelial annotations, the original MACE labels, and SpaceMACE-refined labels. *(Top)* Flow from author annotation to MACE output. *(Bottom)* Comparison of SpaceMACE-S (left) and SpaceMACE (right) outputs relative to the author annotation. The large flow of epithelial cells toward normal reflects the low tumor burden in this dataset, consistent with the precision reduction observed for HCC.

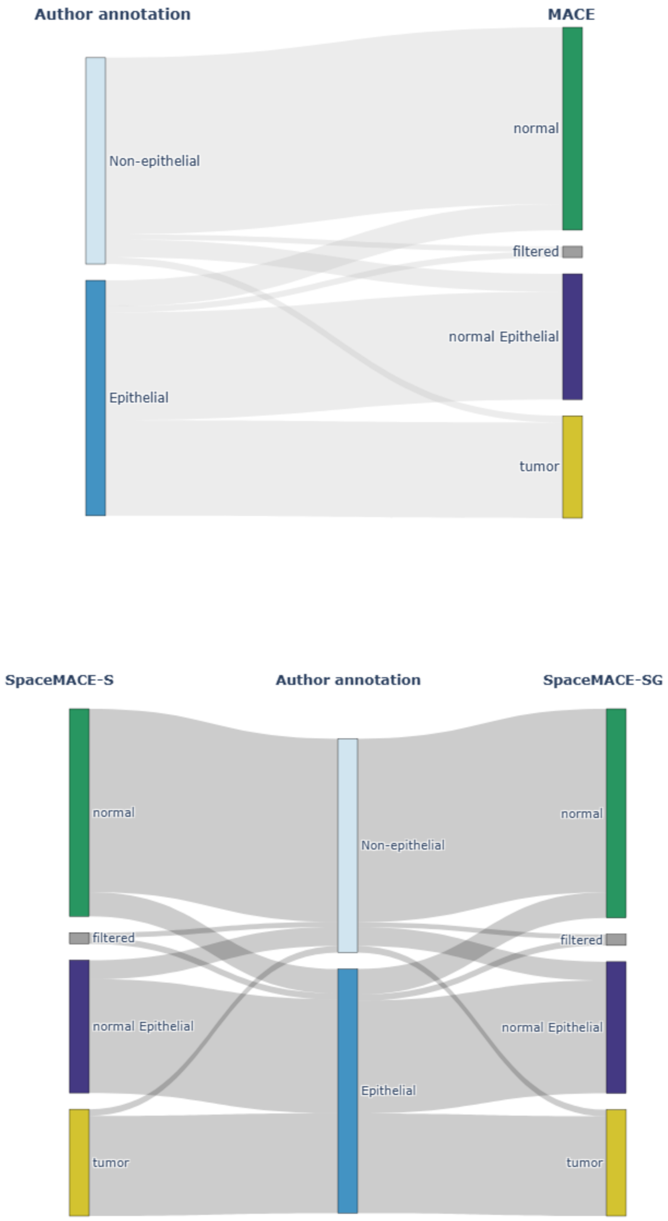


**Suppl. Fig. S10 Detailed label-flow analysis for Stereo-seq OV.** Sankey diagrams show the relationship between author-provided epithelial/non-epithelial annotations, the original MACE labels, and SpaceMACE-refined labels. *(Top)* Flow from author annotation to MACE output. *(Bottom)* Comparison of SpaceMACE-S (left) and SpaceMACE (right) outputs relative to the author annotation. Both SpaceMACE variants show a balance of correct and neutral transitions, consistent with the precision gains observed for OV.

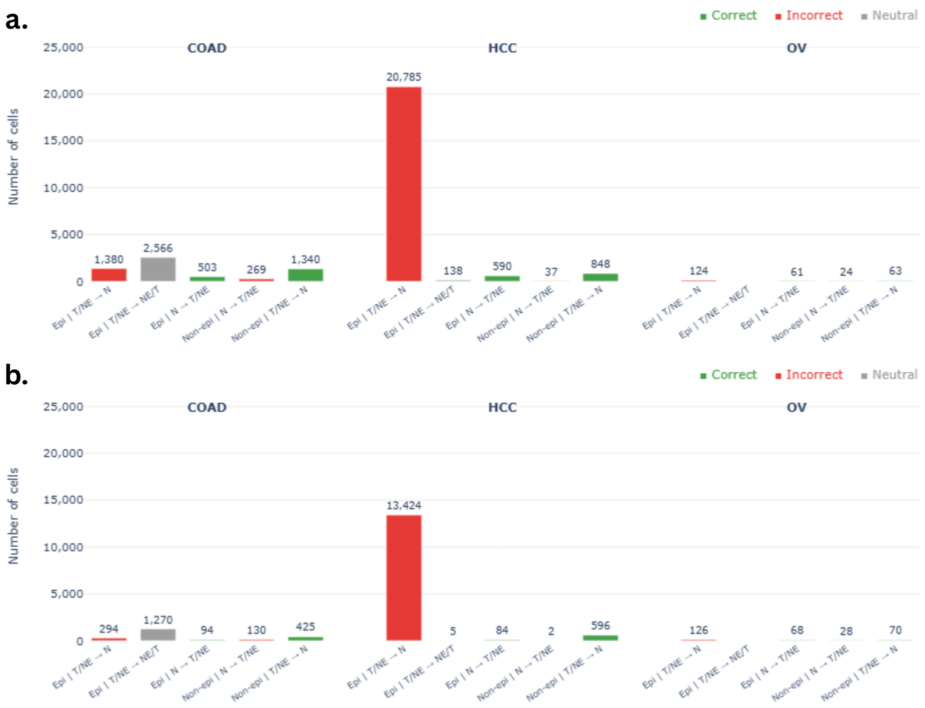


**Suppl. Fig. S11  Cell transition classification across cancer types for SpaceMACE-S and SpaceMACE on VisiumHD (sc) data.** Bar charts showing the number of cells undergoing spatial domain transitions in COAD and HCC, stratified by cell type (epithelial vs. non-epithelial) and transition direction. Transitions are colored by classification outcome: red (Incorrect), green (Correct), gray (Neutral). T, tumor; NE, normal epithelial; N, normal. **(a)** SpaceMACE-S. **(b)** SpaceMACE.

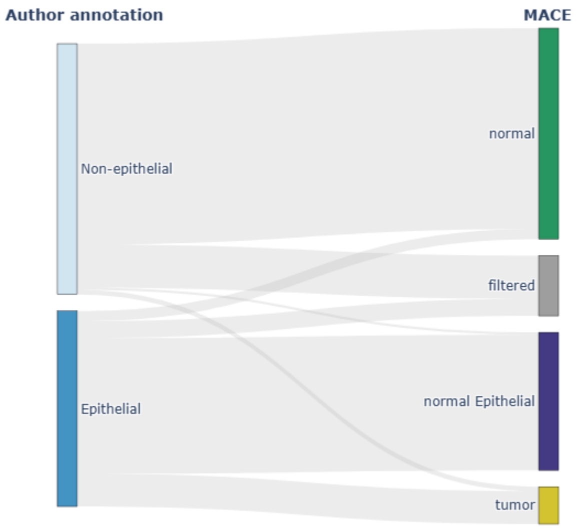


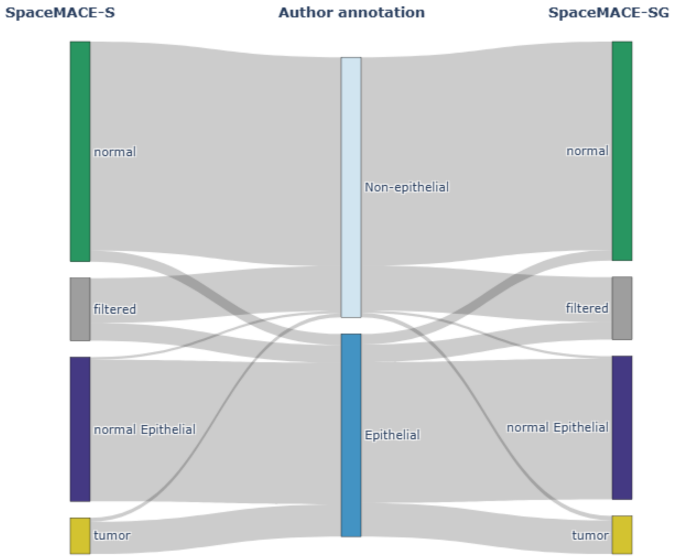


**Suppl. Fig. S12  Detailed label-flow analysis for VisiumHD (sc) COAD.** Sankey diagrams show the relationship between author-provided epithelial/non-epithelial annotations, the original MACE labels, and SpaceMACE-refined labels. *(Top)* Flow from author annotation to MACE output. *(Bottom)* Comparison of SpaceMACE-S (left) and SpaceMACE (right) outputs relative to the author annotation.

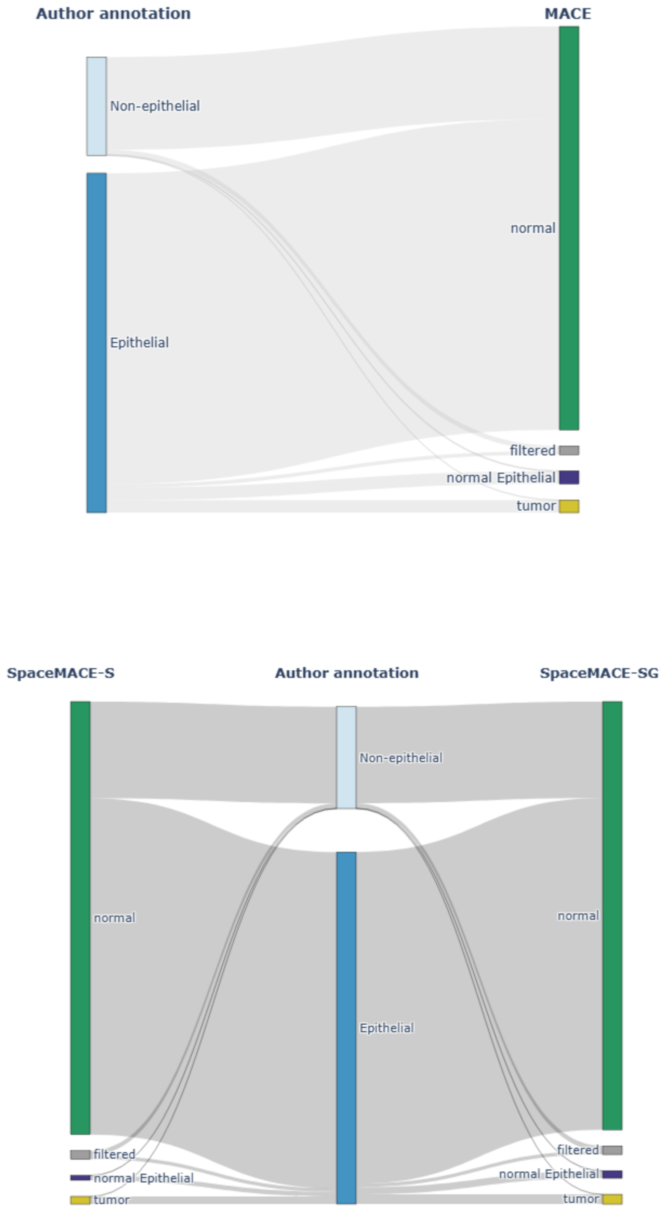


**Suppl. Fig. S13 Detailed label-flow analysis for VisiumHD (sc) HCC.** Sankey diagrams show the relationship between author-provided epithelial/non-epithelial annotations, the original MACE labels, and SpaceMACE-refined labels. *(Top)* Flow from author annotation to MACE output. *(Bottom)* Comparison of SpaceMACE-S (left) and SpaceMACE (right) outputs relative to the author annotation. The dominant flow of epithelial cells toward normal is consistent with the low-tumor-burden limitation observed across HCC datasets.

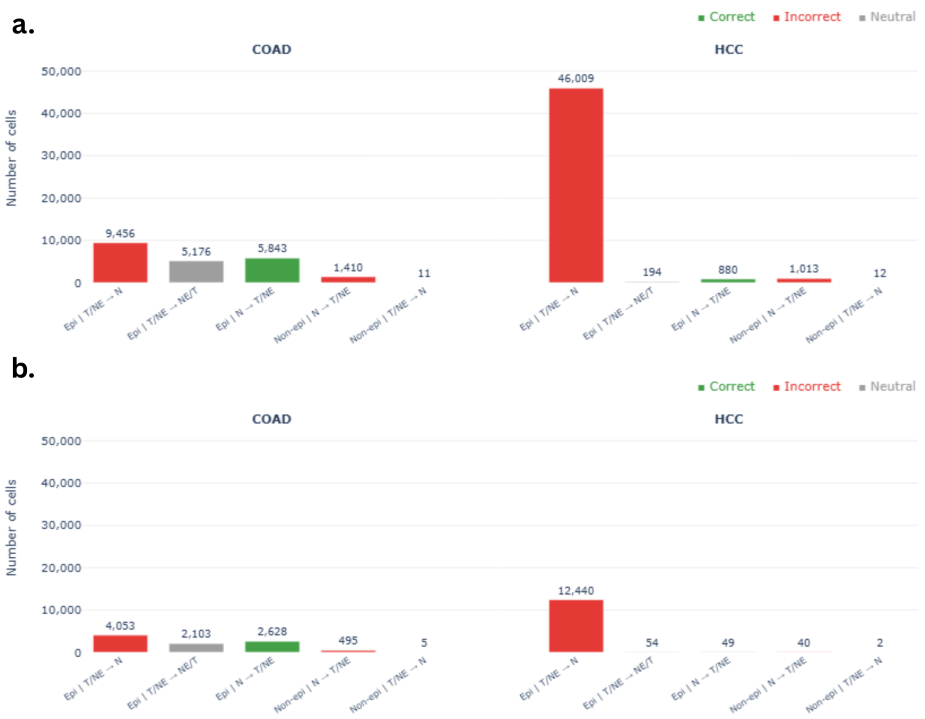


**Suppl. Fig. S14 Cell transition classification across cancer types for SpaceMACE-S and SpaceMACE on Xenium data.** Bar charts showing the number of cells undergoing spatial domain transitions in COAD and HCC, stratified by cell type (epithelial vs. non-epithelial) and transition direction. Transitions are colored by classification outcome: red (Incorrect), green (Correct), gray (Neutral). T, tumor; NE, normal epithelial; N, normal. **(a)** SpaceMACE-S. **(b)** SpaceMACE.

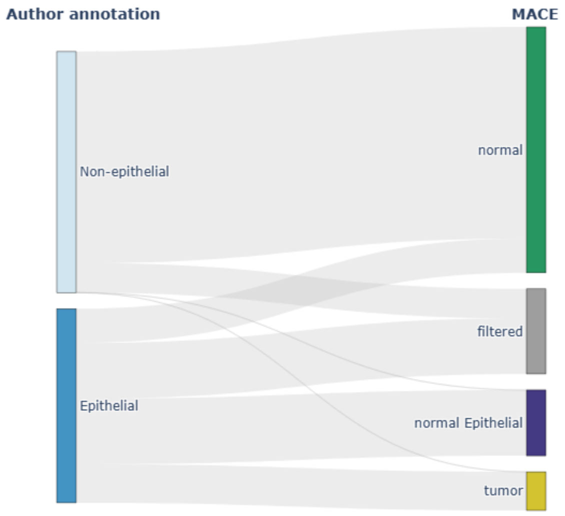


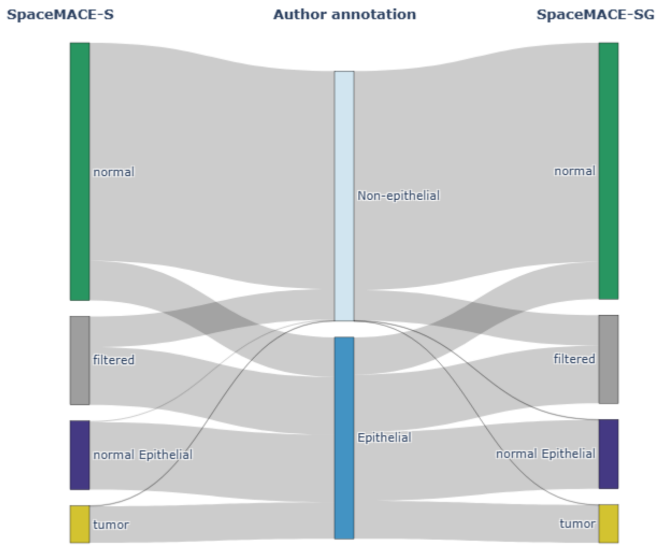


**Suppl. Fig. S15 Detailed label-flow analysis for Xenium COAD.** Sankey diagrams show the relationship between author-provided epithelial/non-epithelial annotations, the original MACE labels, and SpaceMACE-refined labels. *(Top)* Flow from author annotation to MACE output. *(Bottom)* Comparison of SpaceMACE-S (left) and SpaceMACE (right) outputs relative to the author annotation.

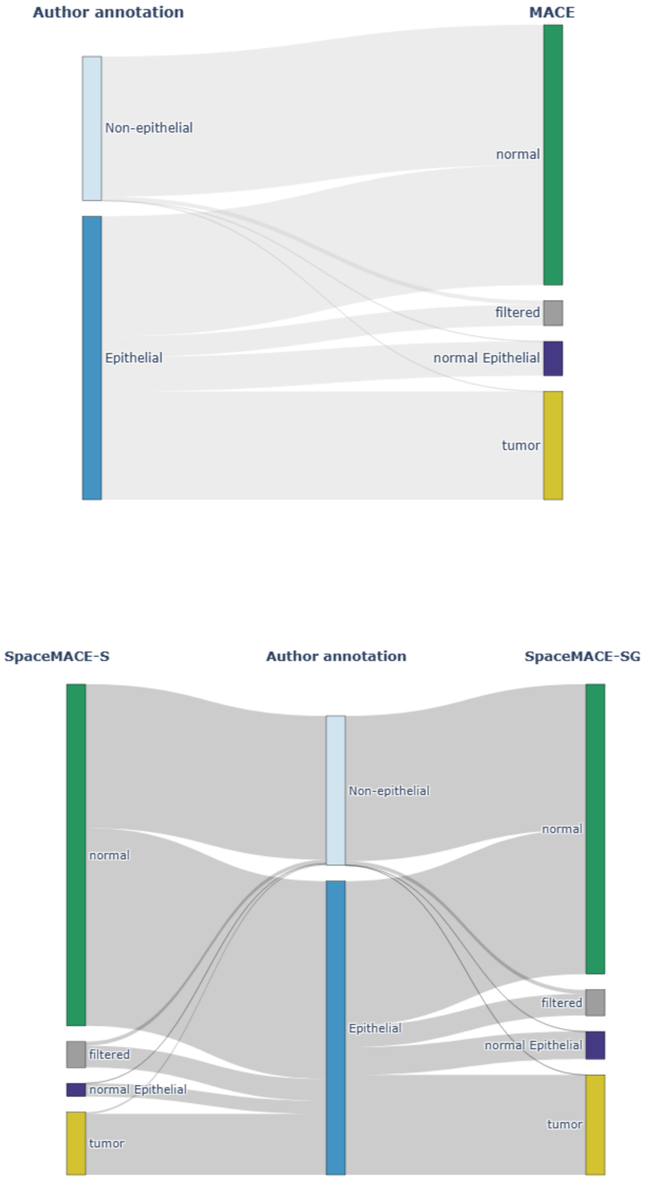


**Suppl. Fig. S16 Detailed label-flow analysis for Xenium HCC.** Sankey diagrams show the relationship between author-provided epithelial/non-epithelial annotations, the original MACE labels, and SpaceMACE-refined labels. *(Top)* Flow from author annotation to MACE output. *(Bottom)* Comparison of SpaceMACE-S (left) and SpaceMACE (right) outputs relative to the author annotation. The predominant reassignment of epithelial cells toward normal reflects the low tumor burden characteristic of HCC datasets.

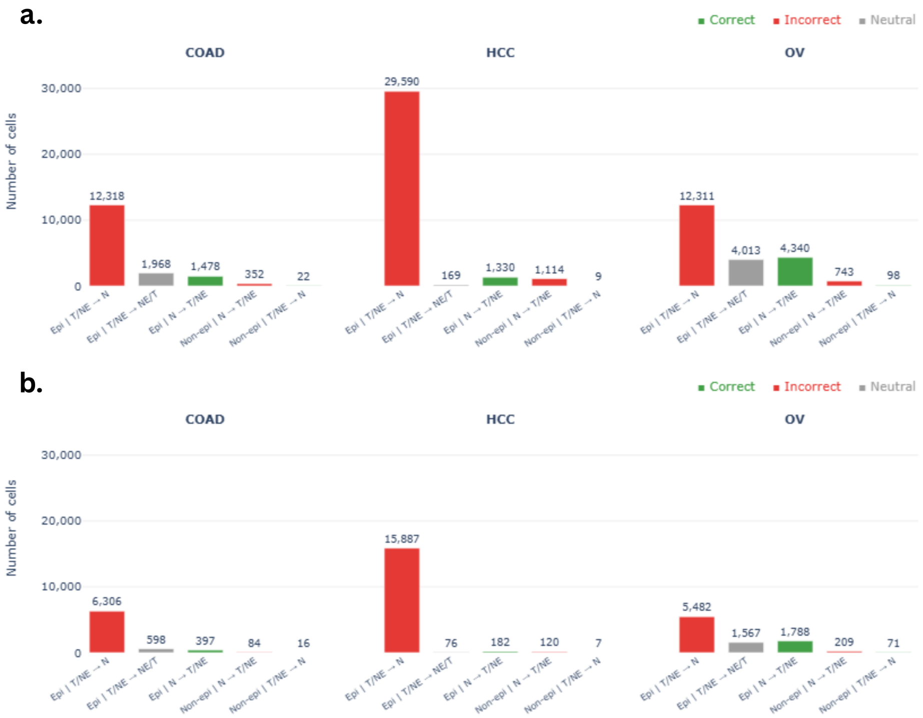


**Suppl. Fig. S17 Cell transition classification across cancer types for SpaceMACE-S and SpaceMACE on CosMx data.** Bar charts showing the number of cells undergoing spatial domain transitions in COAD, HCC, and OV, stratified by cell type (epithelial vs. non-epithelial) and transition direction. Transitions are colored by classification outcome: red (Incorrect), green (Correct), gray (Neutral). T, tumor; NE, normal epithelial; N, normal. **(a)** SpaceMACE-S. **(b)** SpaceMACE.

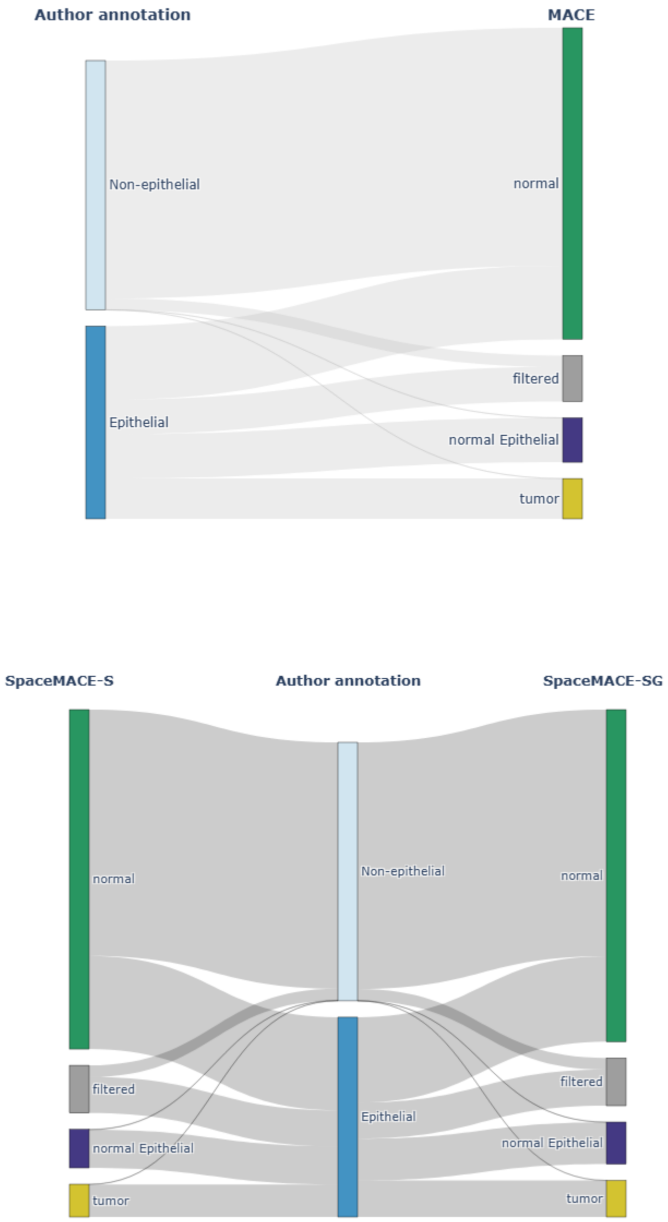


**Suppl. Fig. S18 Detailed label-flow analysis for CosMx COAD.** Sankey diagrams show the relationship between author-provided epithelial/non-epithelial annotations, the original MACE labels, and SpaceMACE-refined labels. *(Top)* Flow from author annotation to MACE output. *(Bottom)* Comparison of SpaceMACE-S (left) and SpaceMACE (right) outputs relative to the author annotation.

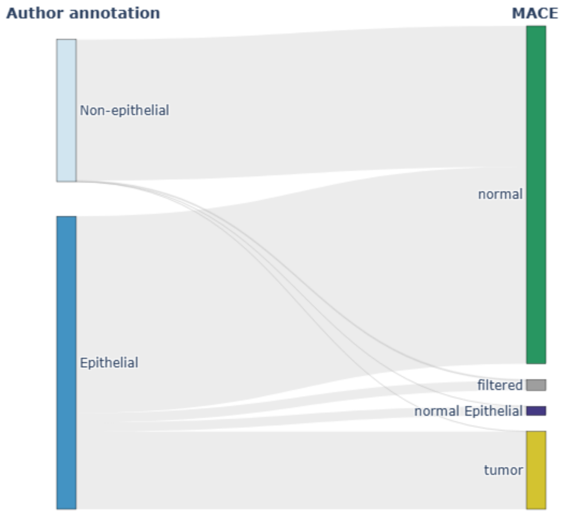


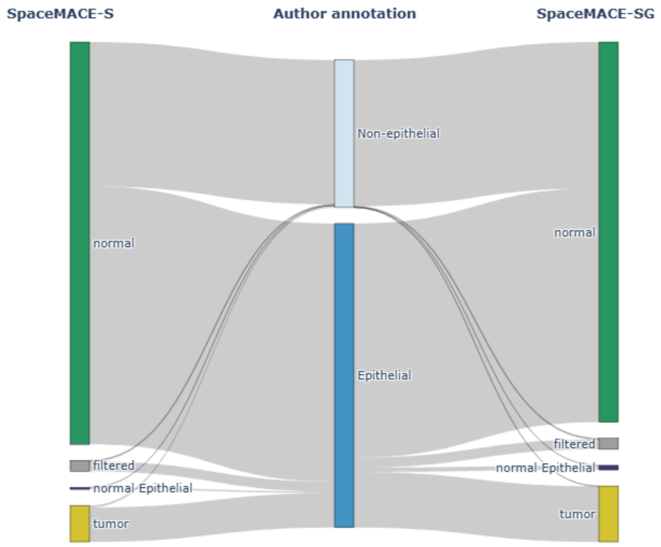


**Suppl. Fig. S19 Detailed label-flow analysis for CosMx HCC.** Sankey diagrams show the relationship between author-provided epithelial/non-epithelial annotations, the original MACE labels, and SpaceMACE-refined labels. *(Top)* Flow from author annotation to MACE output. *(Bottom)* Comparison of SpaceMACE-S (left) and SpaceMACE (right) outputs relative to the author annotation. As with other HCC datasets, the dominant transition reflects mass reassignment of epithelial tumor cells toward normal due to low tumor cell density.

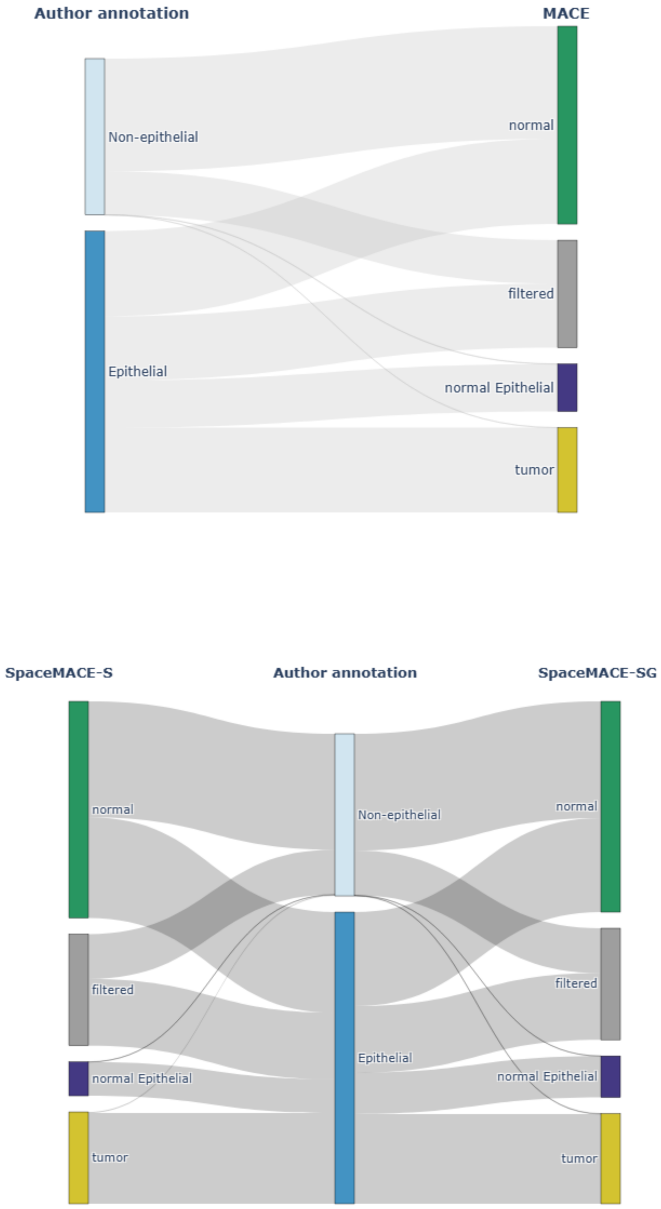


**Suppl. Fig. S20 Detailed label-flow analysis for CosMx OV.** Sankey diagrams show the relationship between author-provided epithelial/non-epithelial annotations, the original MACE labels, and SpaceMACE-refined labels. *(Top)* Flow from author annotation to MACE output. *(Bottom)* Comparison of SpaceMACE-S (left) and SpaceMACE (right) outputs relative to the author annotation. Both variants show a favorable balance of correct over incorrect transitions, consistent with precision gains observed for OV.
